# Microsecond Molecular Dynamics Simulations of Diphtheria Toxin Translocation T-Domain pH-Dependent Unfolding in Solution

**DOI:** 10.1101/572040

**Authors:** Jose C. Flores-Canales, Nikolay A. Simakov, Maria Kurnikova

## Abstract

Diphtheria toxin is a multi-domain protein that invades cells by using their own endocytosis mechanism. In endocytosis, an endosome, a lipid bilayer vesicle, is formed to encapsulate an extracellular molecule. Subsequent acidification of endosome internal solution induces conformational rearrangements and membrane insertion of such encapsulated diphtheria toxin translocation domain (T-domain). In solution at neutral pH, a stand-alone T-domain adopts an all alpha-helical globular structure; however, atomistic details of the pH-dependent conformational changes of the protein are not completely understood. We model structural rearrangements in T-domain in 18 µs long molecular dynamics (MD) simulations of neutral and low pH T-domain models in explicit solvent. At low pH, six histidine residues of the protein were protonated. Two independent MD trajectories resulted in partial protein unfolding at low pH, in which similar regions of the protein conformational subspace were explored. Notably, a pH induced unfolding transition was initiated by partial unfolding of helix TH4 followed by unfolding of helix TH1. Helix TH2 repeatedly unfolds in the low pH T-domain model, which is consequently predicted to be disordered by a consensus of disorder prediction algorithms. Protonation of histidines disrupted a hydrophobic core containing a putative transmembrane helix TH8, which is encircled by hydrophobic surfaces of helices TH3, TH5 and TH9. Afterwards, the low pH T-domain model was reorganized into an ensemble of partially unfolded structures with increased solvent exposure of hydrophobic and charged sites. Thus, MD simulations suggest the destabilizing role of protonation of histidines, in the neutral pH conformation in solution, which may facilitate the initial stages of T-domain membrane binding. The simulation at neutral pH samples conformations in the vicinity of the native structure of the protein. However, significant fluctuations of the protein, including unfolding and refolding of α-helices were observed at these simulation time-scales.

## 1. INTRODUCTION

Diphtheria toxin is a 535 aminoacid protein secreted by pathogenic strains of Corynebacterium diphtheria. Its function is to inhibit protein synthesis of sensitive eukaryotic cells.^1-3^ The toxin consists of a receptor, a translocation, and a catalytic domains with a disulfide bridge between the translocation and catalytic domains. The diphtheria toxin cell insertion proceeds in the following stages:^1^ (i) the receptor domain binds to a cell receptor causing cell absorption of the toxin-receptor complex; (ii) an acidic environment in the endosome interior triggers membrane insertion of the diphtheria toxin translocation (T) domain; (iii) this is followed by the permeation of the attached catalytic domain and its subsequent release into the cytoplasm. Once in the cytoplasm of the target cell the catalytic domain disrupts protein synthesis leading to cell death.^4^ The second stage of this process is of fundamental interest because of the T-domain unassisted membrane insertion,^5^ which also occurs when a T-domain is in the absence of the rest of the protein and of the receptor.6 This contrasts the assisted insertion of membrane proteins and translocation of water soluble proteins by SecY translocon complex,^7^ and makes T-domain a suitable system for pharmaceutical applications.^8,9^ The latter studies have shown that the membrane insertion of the T-domain and delivery of the attached catalytic domain are robust to genetic alterations of the receptor binding domain designed to kill specific cancer cells.^8,9^

Numerous experimental studies were performed to elucidate the structure^10^ and mechanisms^2,3^ of the entire toxin and a stand-alone T-domain. However, atomistic details of the T-domain transformation along the membrane insertion pathway are yet to be established. At neutral pH, T-domain is an all α-helical protein formed with ten helices TH1-TH9 and TH5’. The solvent accessible helices TH1-4 and TH5-7, TH9 encircle a non-polar helix TH8 buried in the protein core (see Figure 1 for illustration of the protein structure).^11^ Lowering pH in solution triggers formation of T-domain intermediate states that brings the system from a soluble to a trans-membrane state, in which the hydrophobic helix hairpin TH8-9 spans the membrane. Possible interfacial and partially inserted states of helices TH1-7 are also indicated via indirect observations.^6,12-15^ Since the interior of an endosome has a pH value close to pH 5^16^ the six histidine residues were implicated in triggering pH-dependent structural changes of T-domain.^17-21^ In addition, pH-dependent local conformational changes of T-domain in solution were indicated via indirect experimental observations.^22,23^ Thus, it is important to understand the role of protonation of histidines in the triggering of conformational changes in solution.

**Figure 1.**
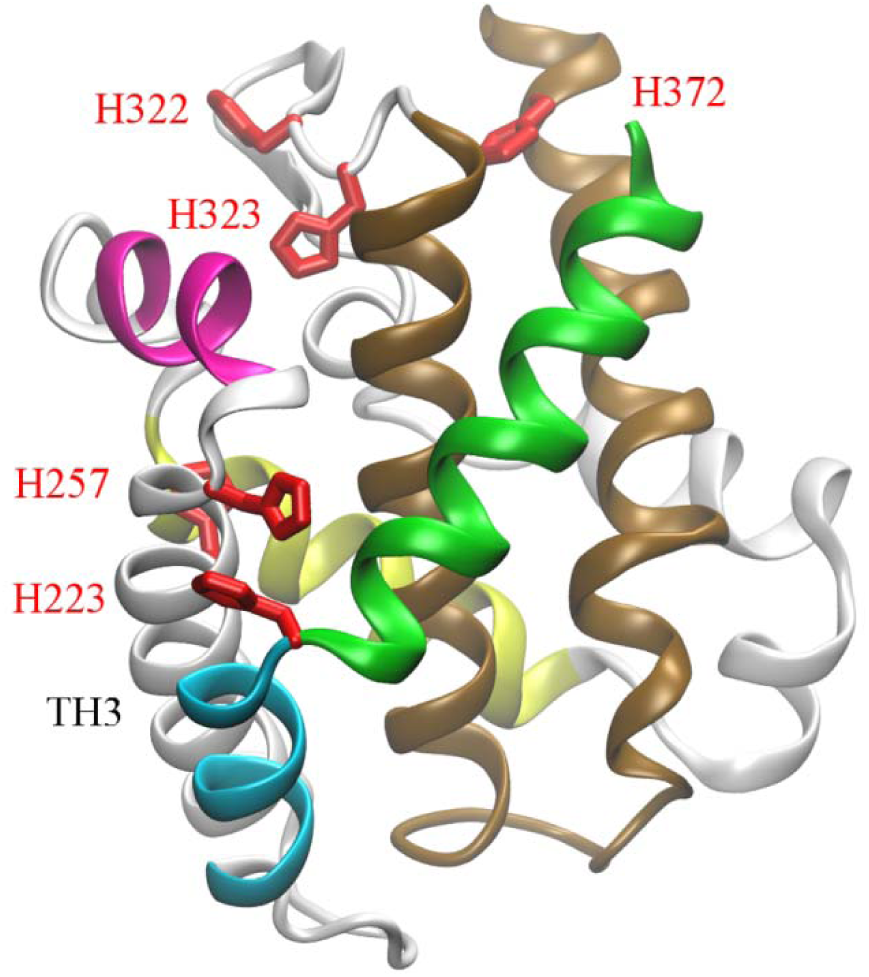
Crystal structure of T-domain extracted from diphtheria toxin monomer structure at neutral pH [PDB ID 1F0L]. Helices TH1 and TH2 are colored green and cyan, respectively. Helices TH4 and TH5 are colored magenta and yellow. Helices TH8 and TH9 are colored brown, the rest of the structure is in grey. Helix TH3 is labeled. Histidine side-chains are shown red in licorice representation and labeled. H251 is buried within helices TH3 and TH5.

We have recently reported on the shifts of T-domain histidine pK_a_s.^24^ using molecular dynamics (MD) free energy calculations (Thermodynamic Integration).^25^ Our results indicated that protonation of certain histidines (e.g. His 257 or His 251) can significantly destabilize T-domain globular structure. Also, our MD simulations suggested that upon protonation of histidines in solution T-domain undergoes partial unfolding resulting in structures in good agreement with spectroscopic experiments.^24^ However, the initial sequence of conformational changes in preparation for subsequent membrane binding has not been characterized.

To gain an insight into structure, diversity and stability of a low pH conformational space of T-domain, we performed extensive 18 microsecond long MD simulations of partial unfolding of T-domain in solution at low and neutral pH. Recent advances in the field of molecular dynamics simulations, such as development of specialized hardware Anton^26,27^ and improved atomistic force-fields^28,29^ make it possible to perform micro- to millisecond-long simulations of protein dynamics in explicit solvent. Atomistic simulations of dynamics of disordered structures and processes of partial unfolding and re-folding in solution are becoming increasingly possible.^30-32^ We demonstrated convincingly that acid driven partial unfolding and refolding of T-domain can occur in simulations on a sub-microsecond time scale.^24^ Significantly larger simulated data sets are needed to fully characterize a partially unfolded state of a protein. At present such fully converged simulation remains an unattainable goal.^30^ Instead, in the present paper we gain insight into T-domain structure in solution at low pH by analyzing similarities and differences in two independent µs long simulations each resulting in protein unfolding and refolding performed at different conditions. The ensemble of these partially unfolded structures has characteristics of a membrane competent state^24^ and remains in a compact conformation.

This paper is further structured into Results and Discussion, and Conclusions sections. The Results and Discussion section first describes simulated models (subsection 1), then it is further structured into three main complementary subsections (subsections 2-4) that together provide comprehensive analysis of the structural changes and dynamics of the protein observed in the simulations. In subsection 2 positional deviation and fluctuation of individual residues and native contacts are discussed in detail; in subsection 3 folding/unfolding of the secondary structure is analyzed using a helicity measure^33^ and its Principal Component Analysis (PCA); and, in subsection 4, we look at the resulting structure and dynamics of the hydrophobic core of the protein and a water exposure of the hydrophobic sites, which are essential characteristics of a protein structure preparation for subsequent membrane association.

## 2. METHOD

### 2.1 Molecular dynamics simulations

The initial T-domain structure was extracted from a high resolution structure (PDB ID code 1F0L).^11^ The simulated protein model contains residues 201-380. Hydrogen atoms were added to both neutral and low pH T-domain structures using tLeap (AMBER 9 package).^34^ A brief description of atomistic models used in MD simulations T1 and T2 follows, a detailed description has been published in a recent report.^24^ N_ε_ atoms were protonated and N_δ_ were not protonated in all histidine side-chains in the neutral pH T-domain structure. The simulation boxes of the neutral and first low pH T-domain model were created by adding 4587 TIP3P explicit water molecules such that the distance between the protein and the simulation box edge was 8.0 Å. The total number of atoms is 16521 atoms in these boxes. In the second low pH T-domain model (T3) reported here, we added 13215 TIP3P explicit water molecules such that the distance between the protein and the edges of the simulation box was 16 Å. This box contains 42405 atoms. Sodium ions were added to the simulation box to neutralize the simulation box. The system was equilibrated with Desmond 2.0 using the ff99SB force field.^28^ The simulation time step was 2 fs and all hydrogen bonds were constrained via SHAKE.^35^ Periodic boundary conditions were set up, cutoff radius was set to 10 Å and electrostatic calculations were performed using Particle Mesh Ewald (PME) method.^36^ The protein was restrained and the solvent was minimized for 200 steps of steepest descent followed by 50 steps of conjugate gradient descent minimization method. Then the solvent was restrained and the protein with backbone atoms restrained was minimized by a total number of 250 steps as explained above. Isotropic NPT ensemble equilibration was performed using MTK barostat^37^ at T = 300 K and pressure of 1 atm with positional restraints applied to heavy atoms (1.0 kcal/mol·Å^2^) over the first 600 ps. Then, the restraint constants were decreased linearly to 0.0 kcal/mol·Å^2^, over the following 400 ps. This was followed by unrestrained NPT equilibration over 8 ns. Production MD simulations were performed on Anton supercomputer.^26^ The ff99SB force field is used and electrostatic interactions were computed by Gaussian Split Ewald method^38^ with a grid size of 64 × 64 × 64 and a cutoff radius of 12.0 Å. The time step was 2 fs and all hydrogen bonds were constrained. Berendsen thermostat and barostat were set to T = 310 K (tau = 1 ps) and pressure of 1 atm (tau = 2 ps), respectively. Atom coordinates were saved every 60 ps. MD simulations were carried out up to 2060 ns, 6723 ns and 9587 ns for the MD runs T1, T2, and T3, respectively.

### 2.2. Analysis

C_α_ root mean squared deviations (RMSD) as a function of time relative to their initial coordinates found in helices identified in the crystal structure. MD structures were aligned to the X-ray structure excluding C_α_ atoms in the tails or those in the loops and tails. The C_α_-RMSD is calculated excluding C_α_ atoms from the tails and loops identified in the crystal structure. Root mean square deviation (RMSD), distances between individual atoms, averaging of protein structures and secondary structure analysis were calculated using the ptraj program available in AmberTools1.4. Solvent accessible surface area (SASA) was determined using a probe radius of 1.4 Å. Molecular figures were prepared using VMD 1.8.7.^39^ Secondary structure assignments are calculated using DSSP.^40^ The cumulative average of the helicity content per helix in Figure 6 is obtained from the DSSP output over the MD trajectories where the window average is 600 ps. We use a measure of native similarity 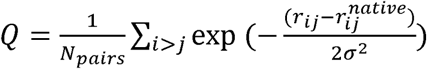, where *N*_*pairs*_ is the number of pairs of native contacts between all Cα atoms in residues 206-375. *r*_*ij*_ and 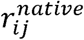 are the distances between Cα atoms in a MD frame and the crystal structure, respectively. σ has a value of 3.0 Å. Contact maps are calculated using the fraction of non-local contacts between the number of pairs *N*_*ij*_ of Cα atoms separated by at least 8 Å and *N*_*max*_ the maximum number of pairs *ij*. Residues containing atoms *i* and *j* are separated by at least 3 residues along the chain.

### 2.3 Principal components Analysis

In this work, given the relatively large size of T-domain (180 residues) and the variety of conformational changes, we choose a metric able to characterize coarse-grained features of the T-domain folded and the partially unfolded ensembles. Based in our previous work on characterization of similarities and differences of different ensembles of unstructured peptides,^33^ we use a helicity measure along the protein sequence for each MD frame (details are described in Ref. 33). To reduce the high-dimensional data obtained, principal component analysis is a suitable and widely implemented method. This method is based on the assumption that important protein motions are described by a linear combination of the main principal components of the covariance matrix; however, non-linear features of MD simulations may not faithfully described by this method.^41^

PCA analysis is carried out by diagonalizing the covariance matrix. Principal components with the largest eigenvalues describe most of the fluctuations in the multidimensional protein space.^42^ This dimensional reduction method can be applied to different metrics i.e.: position of backbone atoms, backbone dihedral angles or helicity measure (triplets).

a. All MD trajectories are stripped of water and ions, followed by translation of the center of mass and orientation alignment relative to the C_α_ atoms of the crystal structure using ptraj program from AmberTools.
b. The set of dimensions *x=* {*x*_1_,*x*_2_,*… x*_*i*_} where *i = 1…p, p* is the number of dimensions, *x*_*i*_ is a vector of size *M*, and *M* is the number of MD frames.
c. A *p* × *p* covariance matrix *R* is defined as follows:

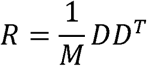

where *D* is a *p* × *M* matrix of elements *D*_*ij*_ *= x*_*ij*_ −< x_*i*_ >, and < x_*i*_ > is the average of *x*_*i*_ over an ensemble of sampled protein conformations.
d. The covariance matrix *R* can be constructed using two different metrics: Helicity measure: A peptide is considered α-helical if at least three sequential residues have their backbone dihedrals (*φ, ψ*) in the α-helical region of the Ramachandran plot, where −100° ≤ *φ* ≤ −30° and −80° ≤ *ψ* ≤ −5°. The helicity for a sequence of three residues is defined as follows:

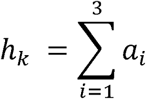

where:

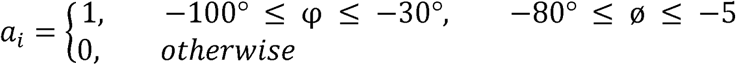

where *i* is the residue index on a kth sequential triplet. The helicity measure is calculated for all consecutive triplet of residues for a protein of *N* residues. This results in a matrix *R* of dimensions *N-2 x N-2*. Another metric is based on the *sin* and *cos* functions of the backbone dihedral angles (φ, ψ) of *N* residues results in a matrix *R* of dimensions *4N x 4N*. In this study PCA analysis is performed over residues 206 – 375.
e. The covariance matrix R is diagonalized to calculate the eigenvalues *λ*^(i)^ and eigenvectors ***v***^(i)^. Those with the largest eigenvalues account for the largest variance in the multidimensional data. Principal components 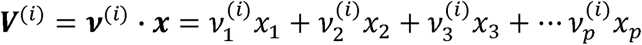. For example, the component 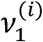 represents the influence of the first triplet comprising residues 206-208 on the ith principal component is represented by 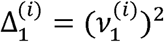. Moreover, the influence of the *ϕ*_1_ backbone dihedral angle on the ith principal component 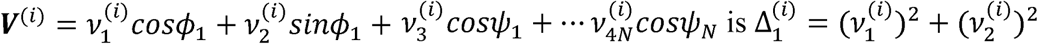.

## 3. RESULTS AND DISCUSSION

### 3.2 Simulated Models of T-domain

The initial coordinates for all T-domain simulations were obtained from the crystal structure of the whole diphtheria toxin protein at neutral pH [PDB ID code 1F0L].^11^ A stand-alone T-domain protein was created by modifying as appropriate the N and C-terminals of T-domain extracted from the full protein. Two models of T-domain were prepared: one with all six histidine residue in their standard protonation states (further referred to as the neutral pH T-domain model); the second one with the histidines in the protonated state (further referred to as the low pH T-domain model). The T-domain protein structure and location of histidine residues are shown in Figure 1. In all simulations all other residues were in their standard ionization states. Three individual MD simulations are presented and analyzed in this paper.

Two of these simulations with neutral and low pH T-domain were performed in water box with dimensions 58 × 59 × 62 Å^3^.^24^ These simulations are referred to as T1 and T2, respectively. The third simulation reported here is a low pH T-domain solvated in a larger water box of the size 78 × 79 × 82 Å^3^, further referred to as T3. This larger system is created in part to assess whether confinement of the protein in a smaller water box (as in T2) affects simulated refolding. For parameters of the simulations please refer to section **2. Method**. 18 µs of atomistic MD simulations of T-domain in explicit water were performed and analyzed in this study.

### 3.2 Simulated Structures and Fluctuations

Root mean squared deviation (RMSD) of C_α_ atoms is a rough measure of structure stability or deformation in the simulations. Figure 2 shows time-evolution of an average RMSD value (top panels, black lines) in each simulation (T1, T2 and T3). RMSD in the neutral pH trajectory T1 (Figure 2A) increased slightly with time to an average value of 1.9 Å, which is somewhat higher than ones typically reported in stable protein MD simulations on nanosecond time-scales. However, similarly high RMSD values were reported, e.g., in a millisecond long simulation of a stable natively folded BPTI protein.^27^ The RMSD value in the range of 2 - 3 Å can be indicative of populating sub-states of the native structure that are inaccessible in shorter simulations. Specifically, such relatively high RMSD may be due to repositioning of the loops, which were confined in the crystal structure because of the crystallographic conditions. This can be demonstrated by leaving out the loops and computing average RMSD of atoms that comprise α-helixes only. Such “truncated” RMSD traces are also shown in Figure 2 in top panels for all three simulations.

**Figure 2.**
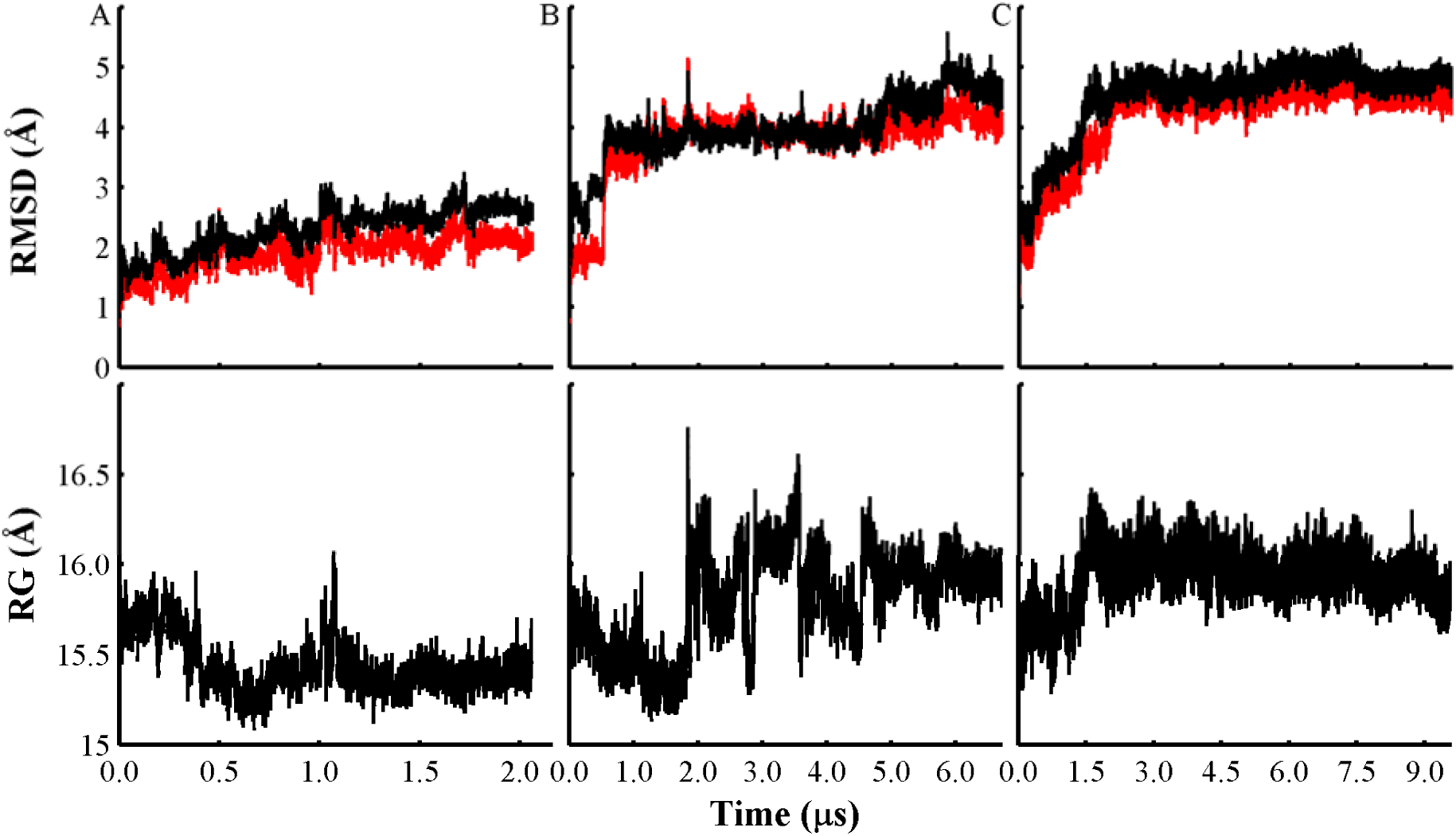
Panels (A), (B), and (C) display C_α_ root mean squared deviation (RMSD) and radius of gyration (RG) calculated from the trajectories T1, T2, and T3, respectively. C_α_-RMSD obtained after C_α_ alignment of MD frames to the initial X-ray structure excluding C_α_ terminal atoms (RMSD computed on residues 206-375 as shown in black lines), and excluding C_α_ atoms in the protein loops and terminals (red lines). Radius of gyration over time for each MD trajectory is shown in lower panels (C_α_ atoms of the residues 206-375).

For the neutral pH simulation (T1) the truncated RMSD is noticeably lower than the full RMSD indicating that the structure is stable and maintains its native fold. This observation is further corroborated by decrease in the radius of gyration in T1 (see Figure 2A lower panel), which is a measure of the overall compactness of the protein structure. C_α_-RMSD traces for both low pH T-domain trajectories T2 and T3 show an abrupt increase to approximately 3 Å in the first microsecond of the simulation, followed by gradual increase to ca. 5Å over the last microsecond (see, Figures 2B, C, top panels). However, the dynamics of the RMSD evolution towards higher values is different in these trajectories. In T2 the initial deformation of the protein is mainly due to the loop deformation (compare black and red lines in Figure 2B in the first 0.5 µs), followed by a sharp increase in the “truncated” RMSD (see ibid.), indicating a change within the helical structure of the protein. In T3, the RMSD changes gradually over a similar period of time (see Figure 2C). Both low pH trajectories exhibit increase in the radius of gyration (Figures 2B, C, lower panels) indicating unfolding. The radius of gyration in T2 decreases initially in a manner similar to T1, then increases abruptly, fluctuates further on and stabilizes at a slightly elevated value at the very end of the trajectory. In T3 the increase is sharp over the initial 1.5 µs, but it remains stable, and decreases slightly by the end of the trajectory, which may be indicative of formation of a new structure

To determine which residues contribute most to the observed average RMSD increase discussed above, it is instructive to plot RMSD values per C_α_ atom averaged over the last stable trajectory segments, further termed at-RMSD (see Figure 3A). In Figures 3B-D representative protein structures from each trajectory are colored according to the corresponding at-RMSD values in Figure 3A. In T1 at-RMSDs are generally small, with the largest Cα atom displacements in two inter-helical loops TH7-8 and TH8-9. This is a typical result for a stably folded protein in a simulation. In contrast, in T2 and T3 a significant fraction of at-RMSDs have large values due to overall structural changes. However, the preserved protein core is clearly indicated by the low at-RMSDs, which comprises helixes TH3, the TH5 C-terminus and TH8. The largest deformation of the structure in T2 (C_α_-RMSD > 4.0 Å) occurs in helices TH1-2, in the N-terminus and in the inter-helical loops TH7-8, TH8-9. In addition to these protein regions, T3 has C_α_-RMSD values larger than 4.0 Å in the TH9 C-terminus.

**Figure 3.**
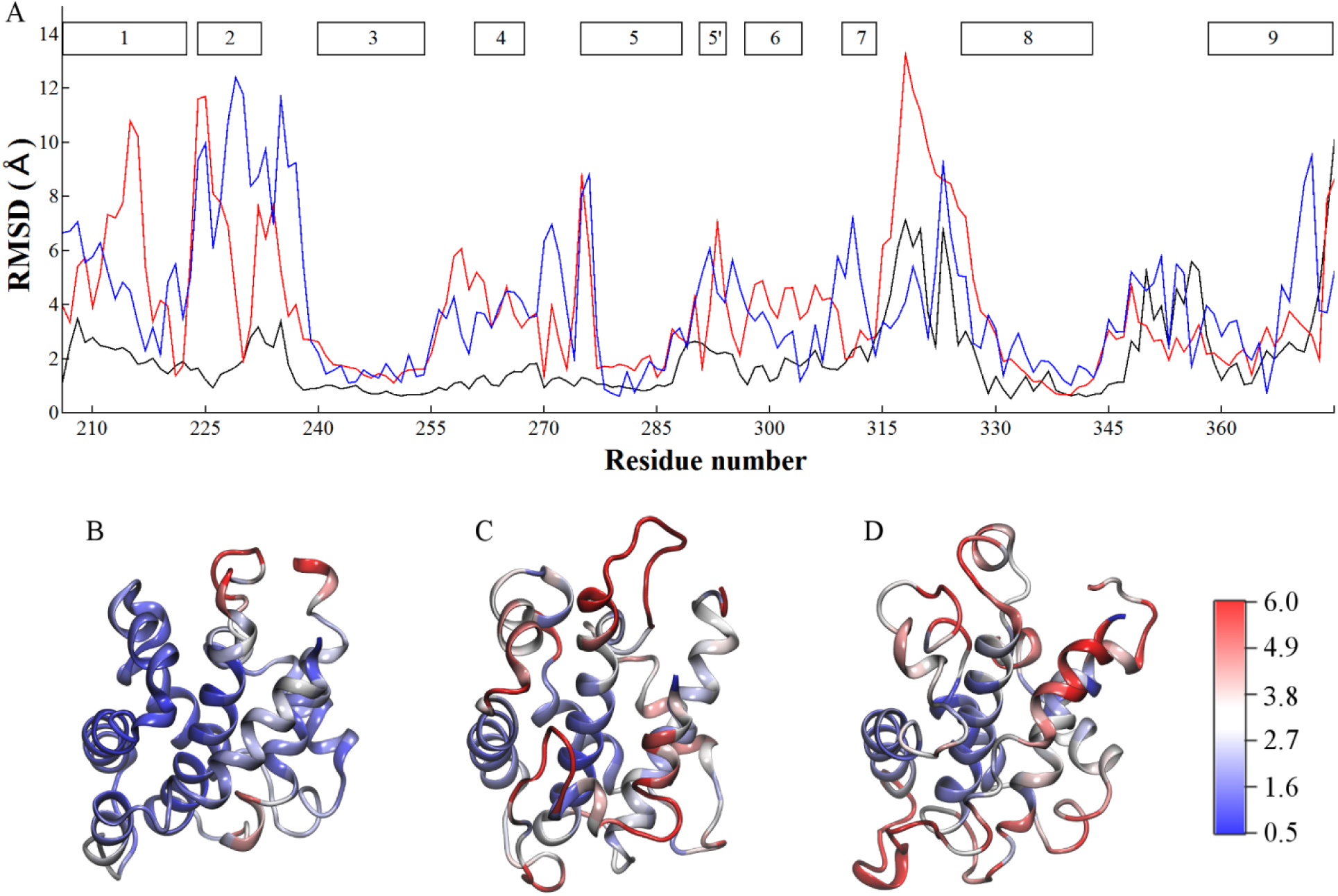
C_α_-Root mean square deviation (RMSD) versus residue number calculated after translation and best alignment of each MD frame relative to the C_α_ atom coordinates of the crystal structure (residues 206 to 375). (A) C_α_-RMSD traces obtained from the last 1 µs of trajectories T1 (black line), T2 (red line), and the last 2 µs of T3 (blue line). In addition, helice TH1-9 and TH5’ are indicated by grey filled rectangles (top of the C_α_-RMSD plot). Ribbon representation of representative structures colored according to their C_α_-RMSD, see the color bar. Structures are obtained from: (B) The last 1 µs of trajectory T1. (C) The last 1 µs of trajectory T2. (D) The last 2 µs of trajectory T3. A representative structure has the lowest C_α_-RMSD relative to the average structure over the last segment considered of each trajectory.

Mobility of atoms with respect to their average position (determined here as root mean squared fluctuations (RMSF) of C_α_ atoms in the final stable segments of the trajectories, see Figure 4) is a useful characteristic for distinguishing unstructured and poorly structured (floppy) from folded conformations. Overall in both low pH simulations the protein adopts a fairly stable structure characterized by low RMSF of the majority of the residues, comparable to the RMSF of the natively folded structure simulation T1. The main difference in local mobility of residues in T2 and T3 simulations arises from the inter-helical loop TH7-8. In T2 its RMSF values reach 3.0 Å as a result of an unstructured conformation (Figure 4A, C). In T3 RMSFs are lower than 2 Å in this region, which is partially folded in a helical structure. Note that protonated H322 and H323 are located in this loop. SASA calculations show that side-chains of H322-H323 are largely solvated due to their unstructured conformation in all MD trajectories except in the trajectory T1 where H322 has smaller solvent exposure. The inter-helical loop TH8-9 (Figure 4A, D) is more mobile in T3 than in T2.

**Figure 4.**
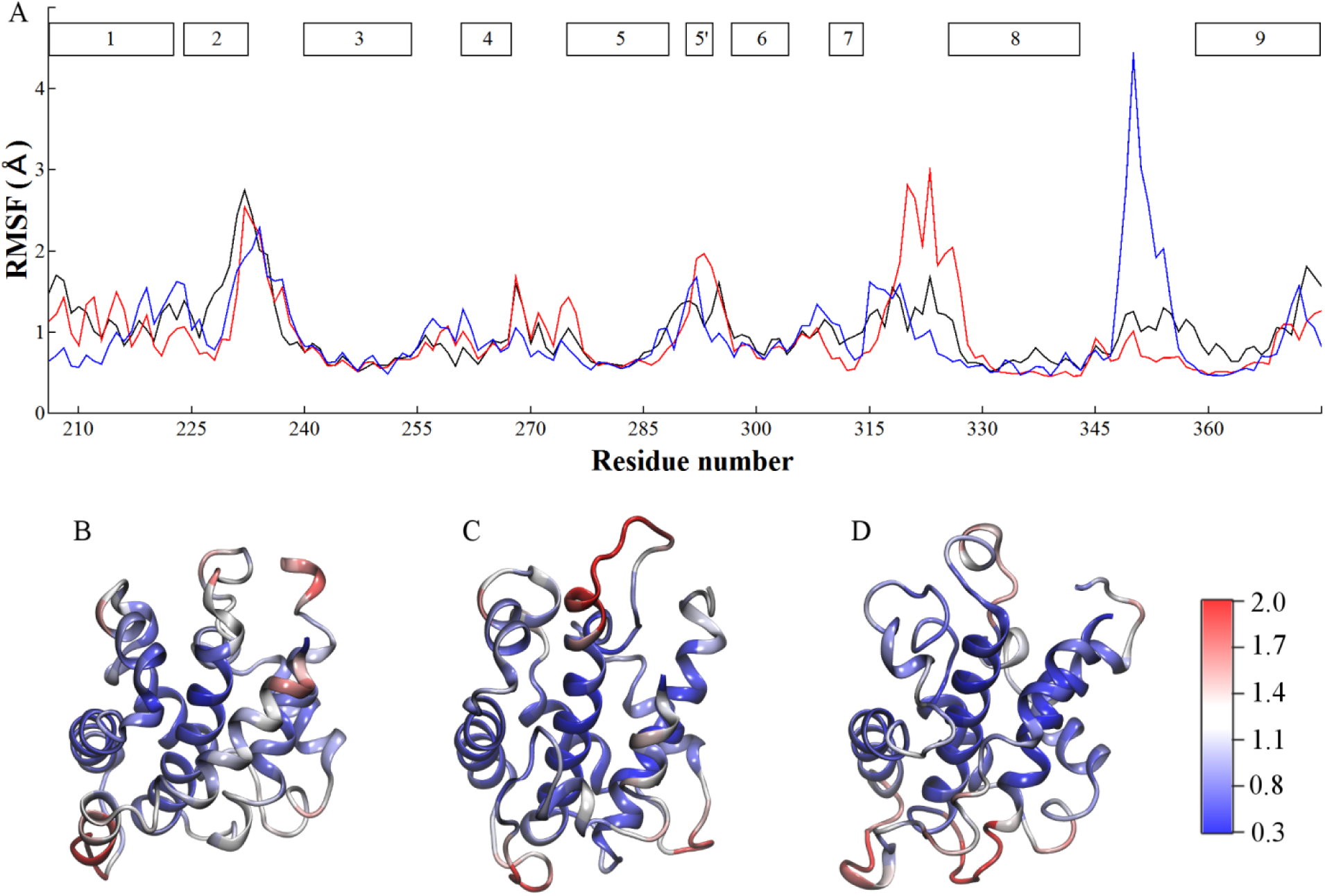
C_α_-Root mean square fluctuations (RMSF) versus residue number obtained after translation and best alignment of MD structures relative to the C_α_ atom coordinates of each representative structure (residues 206-375). (A) C_α_-RMSF traces obtained from the last 1µs segments of trajectory T1 (black lines), T2 (red lines), and the last 2 µs of T3 (blue lines). Helices TH1-9 and TH5’ are represented by filled grey rectangles (top of C_α_-RMSF plot). Ribbon representation of representative structures colored according to their C_α_-RMSF values, see the color bar. Structures are obtained from: (B) The last 1 µs of trajectory T1. (C) The last 1 µs of trajectory T2. (D) The last 2 µs of trajectory T3. A representative structure is the closer to the average structure calculated over the last segment of each trajectory.

Disruption of a native structure is best characterized with an order parameter *Q*, whose value reflects on a number of intact native contacts in the structure (see section **2. Method** for definition of *Q*). In Figure 5, a probability density of the order parameter *Q* is shown for all trajectories. T1 shows two peaks at *Q* = 0.91 and *Q* = 0.86. The second peak is populated by the structures in which one salt bridge R210-E362 is disrupted. Both low pH T-domain trajectories show a similar peak around *Q* = 0.79 populated during the first 0.5 µs of the trajectories (and before unfolding of the helices TH1, TH2, see further analysis in subsection **3.3 Secondary Structure Dynamics**). After unfolding of these helices, T2 and T3 populate peaks centered at *Q* = 0.67 and *Q* = 0.73, respectively. Further refolding of T-domain in T2 results in structures with *Q* = 0.61. An effective number of lost native contacts in T2 is higher than that of T3 trajectory, indicating that the smaller amount of solvating water and consequently smaller simulation box were not severe limiting factors for structural changes of the protein during these simulations. A more detailed analysis of the individual residue-residue contacts lost and formed while the protein underwent structural reformation can be done via analyzing two-dimensional contact maps. These are presented in Supporting Information in Figures S1A-C for all trajectories (see also description of these calculations ibid.). In both T2 and T3 simulations, the residues of the helix TH1 form non-native contacts with residues in the inter-helical loop TH8-9 and in the TH9 N-terminus (Figure S1B, C). In T2, helix TH1 also loses native contacts with residues in the inter-helical loop TH3-4 and TH8 (Figure S1B). In both T2 and T3 helixes TH2 and TH3 lose a number of native contacts (compare to T1 in Figure S1A). Finally, in both T2 and T3 helix TH4 loses contacts of the residue E362 (TH4) with the residues E327 and Q331 in TH8.

**Figure 5.**
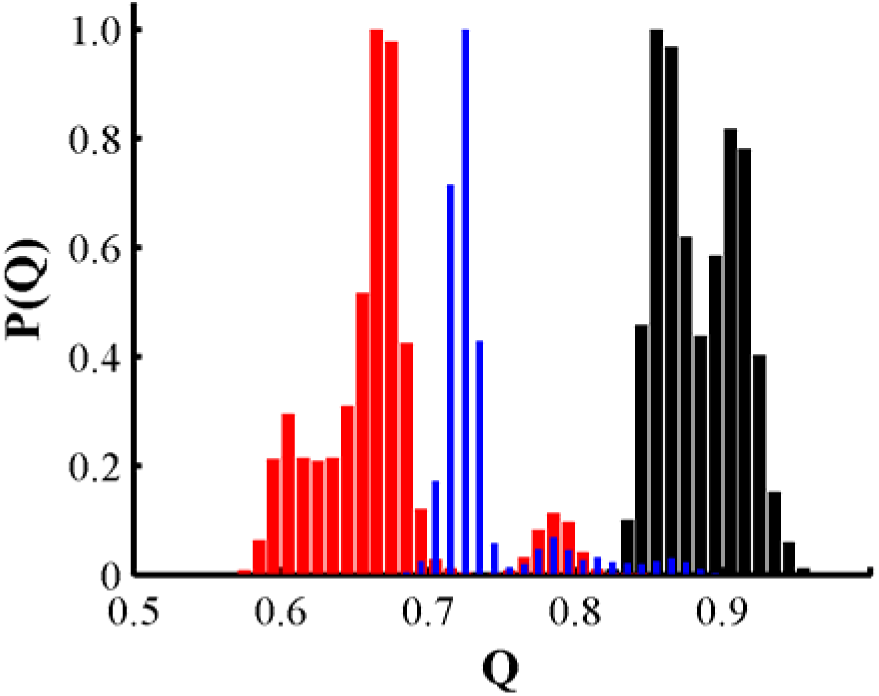
Probability density *P*(*Q*) of native contacts *Q* calculated over C_α_ atoms of residues 206-375. *P*(*Q*) is calculated over trajectory T1 (black line), T2 (red line), and T3 (blue line). A value of *Q* = 1 represents the crystal structure and *Q* = 0 represents an unfolded state.

As indicated by the contact map plots in Figure S1 and at-RMSD plots in Figure 3, helices TH1 and TH2 undergo backbone conformational changes in both low pH trajectories T2 and T3 (structures shown in supporting Figure S2). Protonation-dependent transition of the helix TH1 is initiated by a drastic change of the backbone dihedral angles of K216 in both low pH protein trajectories (Figure S3). In T2 unfolding of TH1 is substantial, characterized by the breaking of inter-helical salt bridges K216-E259, K212-E326, and K212-E327 (Figure S4). In T3 there is a disruption of the first two salt-bridges but the third salt-bridge remains intact. Following this partial unfolding TH1 adopts a helix-kink-helix structural motif in T2 (Figure S2 and Ref. 24). Helix TH2 undergoes a helix to coil transformation in both low pH MD simulations. Such similar behavior in two independent microsecond long simulations highlights the importance of the finely tuned electrostatic interactions in the protein N-terminus disrupted by protonated H223 and H257. Protonation of histidines induces disruption of a structure stabilizing hydrogen bond network (H257-S219 and H257-E259) in both trajectories T2 and T3. Furthermore, conformational changes of helices TH1 and TH2 facilitate solvent exposure and separation of the histidine side-chains. Their solvent accessible surface areas are 100 Å^2^ (H223) and 45 Å^2^ (H257) after the structural transitions in both low pH trajectories, while they are 28.2 Å^2^ (H223) and 17.2 Å^2^ (H257) in T1. Upon transition, the side-chain of H257 is stabilized by a hydrogen bond with the backbone oxygen of A254 in both T2 and T3. In addition, the C-terminus of TH9 is unfolded in both simulations T2 and T3. Finally, an α-helical turn in TH8 C-terminus and TH9 N-terminus is formed in all MD simulations.

Final structures in T2 and T3 show structural difference in the inter-helical loop connecting helices TH7-8 in which final T2 structure exhibits an extended loop conformation and TH7 adopts a 3_10_ secondary structure (after unwinding of residues 313-314). Instead, T3 representative final structure shows a short α-helix (residues 318-321) and a 3_10_ turn (residues 318-320) in the loop between helices TH7 and TH8. Moreover, this loop is stabilized by docking of the protein C-terminal, which is not observed in T2. This is characterized by the stable side-chain distance of Y380 to H323 and formation of a salt-bridge between D377 - E326 along trajectory T3 in contrast to trajectory T2. Correspondingly, the short helix TH7 adopts a 3_10_ secondary structure in both low pH T-domain representative structures (backbone hydrogen bonds are disrupted in residue 313-314). T3 structure shows partial refolding of TH2 into a 3_10_ turn (residues 229-231).

### 3.3 Secondary Structure Dynamics

As shown in the previous section the structure of the T-domain at neutral pH (simulation T1) remained mainly unchanged and was stable in a two microsecond long simulation. An overall shift of residue positions from their initial states, as well as their mobility remains low for the duration of this fairly long simulation. However its secondary structure turns out to be rather dynamic on these time-scales. T-domain is an α-helical protein, and therefore its secondary structure folding and unfolding dynamics is well characterized by monitoring to what extend each α-helix maintains its helical form. The helicity content is calculated as a time-averaged fraction of helical residues that comprise an α-helix identified in the crystal structure (Figure 6). It is somewhat unexpected that the secondary structure of a natively folded stable T-domain (simulation T1) undergoes such significant dynamic transformations in the course of the simulation as shown in Figure 6A. Only three helixes TH3, TH6 and TH8 remain fully helical for 2 µs of the simulation. Helixes TH1 and TH2 are both dynamic, *i.e.* they fluctuate to unfolded states at this time-scale (note the overall structure of the protein remains in the natively folded state, see *e.g.* Figure 5). The most common transition is alternating between α-helix and 3_10_ helix forms (also shown in Figure 6 in red lines), see e.g. TH4, TH7 and TH9 in Figure 6A. A noticeable degree of unfolding into coil structures and refolding is also observed in these helixes. Unlike T1, trajectories T2 and T3 are of a protein that undergoes significant structural transformation in the course of the simulations including irreversible unfolding of some helixes (see Figure 6B, C). Still, there are apparent similarities in the secondary structure dynamics of all three trajectories. In all trajectories helixes TH3, TH6 and TH8 preserve their α-helical content. Early in all trajectories, short helix TH5’ and TH9 C-terminus unfold significantly. Helix TH1 begins to unfold at ca. 500 ns and ca. 2000 ns in trajectories T2 and T3, respectively. Complete unfolding of TH2 occurs approximately at 1400 ns in both T2 and T3. It is possible that the dynamic behavior of TH2 in trajectory T1 is indicative of its propensity to undergo unfolding upon HIS protonation (observed in T2 and T3). Moreover, according to a protein structure disorder prediction meta server^44^ the sequence comprising helix TH2, *i.e.* residues 226-233, is disordered. The meta server employed eight sequence-based disorder predictors, of which at least six predictors qualified this sequence as intrinsically disordered (with disorder prediction consensus larger than 0.8). The TH4 C-terminus unfolds early at 160 ns in T2 and refolds at 1420 ns. In T3, it unfolds at 1530 ns. Despite differences in unfolding behavior of TH4, its unfolding precedes structural changes of TH1 in both low pH T-domain trajectories. Also, TH7 transitions to a 3_10_ conformation at similar times (4630 ns and 5090 ns) in T2 and T3. Conversely, the TH5 N-terminus undergoes a slow transition after 5730 ns in T2 but a rapid one at 1180 ns in T3. (Note that H251 is located in TH3 and within 8 Å of N-terminal residues 275-279 of TH5).

**Figure 6.**
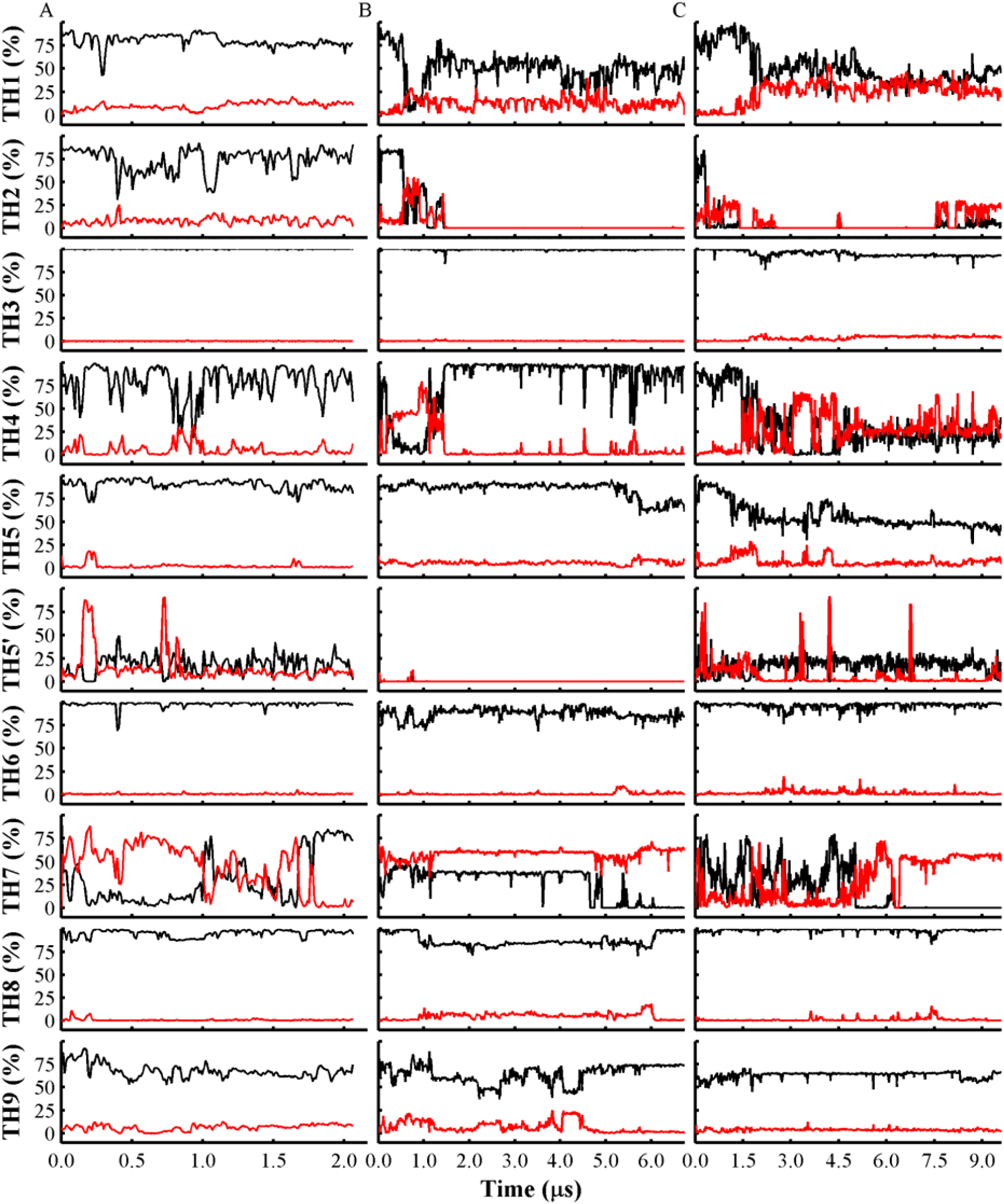
Secondary structure of helices identified in the T-domain crystal structure. (A) Secondary structure along the trajectory T1. (B) Secondary structure along trajectory T2. (C) Secondary structure over the course of trajectory T3. Alpha-helical content (black lines) and 310 content (red lines) are plotted for each alpha-helix identified in the crystal structure. Analysis was performed with DSSP program and cumulative average was calculated with an averaging window of 1.2 ns.

Both similarities and differences are observed in the unfolding paths and the resulting protein structures in trajectories T2 and T3 (see Figure 5). These are difficult to characterize, especially, given an incomplete and non-stationary sampling of the protein conformational space. Yet a joint analysis of the available trajectories is useful for inferring persistent properties of the underlying structural ensembles. To achieve that, a principal component analysis of the joint simulated structural ensembles is designed here. This consists on the projection of trajectories onto the two lowest principal components derived using only stationary basins of folded and refolded structures. The metric used to characterize the structural order is a local helicity measure along the protein sequence introduced in a study of unstructured peptides by Speranskiy and Kurnikova.^33^ The measure is defined for triplets of consecutive residues that is sufficient to determine whether the secondary structure is locally α-helical (a detailed description is provided in **Method** section). Principal components are initially determined using a joint dataset (T1, T2) that contains samples from the last stable segments of simulations T1 and T2 (1 µs each), i.e. using samples corresponding to quasi-equilibrium trajectories of a folded and a refolded protein. By design, first several principal components expose the major structural differences between these two protein states. Indeed, Figure 7A shows that the first two principal components obtained from the first dataset differentiate a folded (T1) and a partially refolded ensemble (T2). The projection of the last 1 µs segment of T3 (also a sample from a quasi-equilibrium trajectory after protein unfolding and reorganization) partially overlaps conformational subspace of T2, (also shown in Figure 7A). This illustrates structural similarities in the ensembles of partially refolded trajectories from two independent simulations. Furthermore, projection of the entire trajectories T2 and T3 on the same PCA vectors (Figure 7C) shows that the conformational subspace covered by trajectory T2 contains that of trajectory T3. Projection of the whole T1 onto the same subspace shows only small variation near its initial folded conformation.

**Figure 7.**
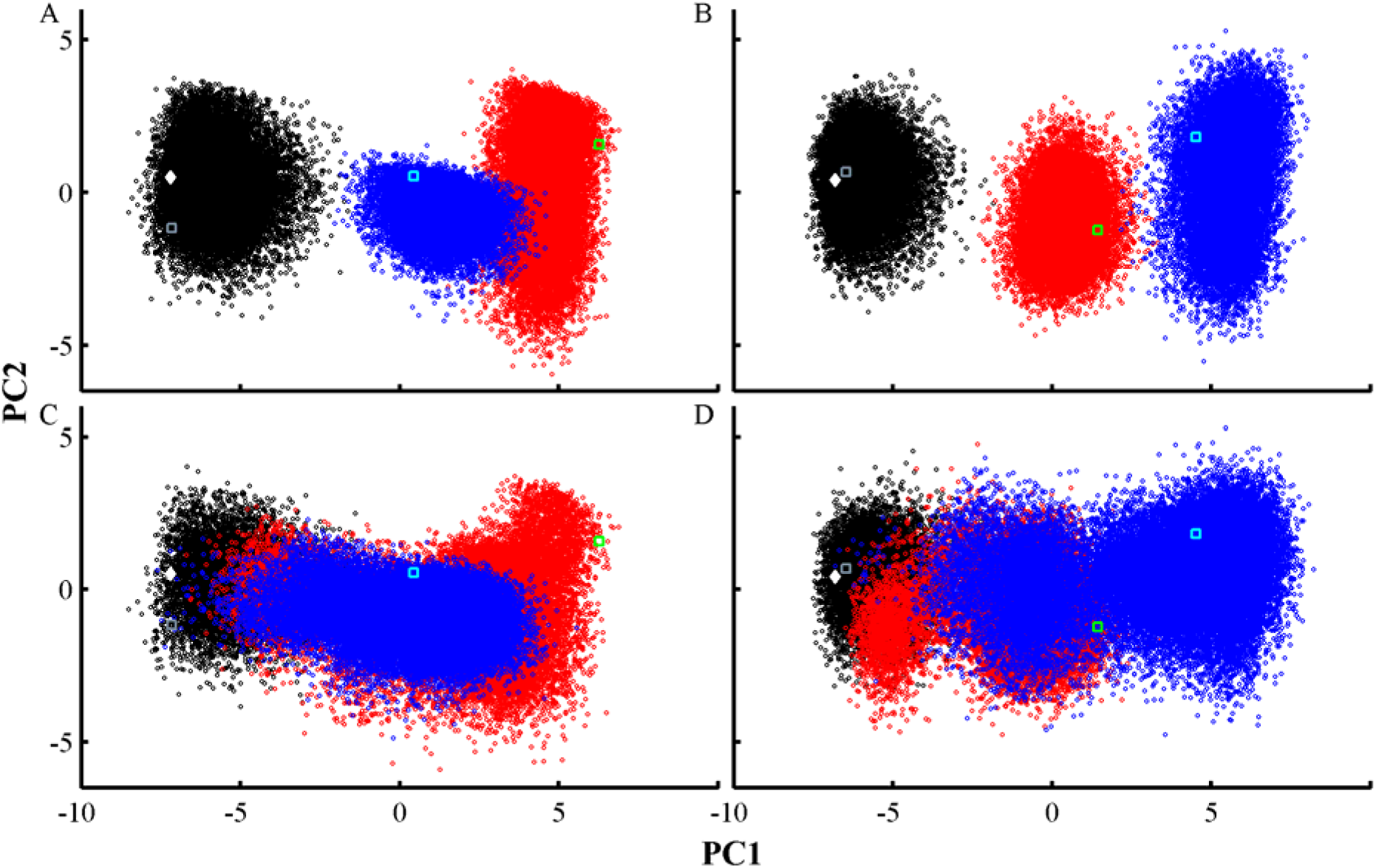
Projection of MD frames on the two main principal components calculated from two datasets comprising the last 1 µs segments of trajectories T1 (black circles), T2 (red circles), and T3 (blue circles). (A) Two-dimensional projection of the last 1 µs of all trajectories on principal components obtained from a dataset containing the last 1 µs of (T1, T2). (B) Two-dimensional projection of the last 1 µs of all trajectories on principal components obtained from a dataset containing the last 1 µs of (T1, T3.) (C) All trajectories are projected on principal components obtained from a dataset containing the last 1 µs of (T1, T2). (D) All trajectories are projected on principal components obtained from a dataset containing the last 1 µs of (T1, T3). Principal component analysis (PCA) was carried out on the triplet measure for each MD frame (residues 206-375). Projection of the X-ray structure is shown in filled white diamond. Final MD snapshots of trajectories T1, T2, and T3 are shown in grey, cyan, and green empty squares, respectively.

In Figure 7B the same procedures are applied to the dataset formed first by joining the last equilibrium segments of trajectories T1 and T3. The samples from the last 1µs of trajectory T2 is then projected onto the two lowest PCA components of this (T1, T3) dataset. In this case the first two principal components of the second dataset show no overlap of the last MD segments of T2 and T3, however projection of the complete MD trajectories on these principal components (Figure 7D) shows, once again, a partial overlap of structural ensembles of T2 and T3. MD frames from T2 and T3 that sample the same region of the projected conformational space correspond to the trajectory segment after the unfolding of TH1 at ca. 0.5 µs of T2 and to the first 5 µs of T3. Thus, the final segment of T3 samples a new set of conformations that are structured differently from ensemble in T2. The first principal component (PC1) contributes ca. 44 – 47 % of the variance in both datasets, see supplemental Figure S5. Furthermore, triplets containing residue H257 have the largest influence on PC1 in both datasets, shown in Figure S6. On the other hand, the second principal component (PC2) is influenced by triplets in helices TH6 or TH7 and by triplets close to residue H223 (residues 218-220 and residues 224-226 on each dataset, respectively).

The helicity measure used above is one way to characterize the secondary structure of a protein. Another structural measure commonly used to characterize subtle differences in protein conformations^43^ is via individual values of the backbone dihedral angles (see **Method** for description). Such measure reports on a slightly different set of structural descriptors and is more sensitive to the specific local conformation of the backbone. For completeness of the structural analysis PCA was also performed in the manner described above for the backbone dihedral angle measure. The results are presented in the Supporting Information.

### 3.4 Protein Hydrophobic Core and Surfaces

Preserving a hydrophobic core and formation of a balanced water exposed surfaces is one of a signature properties of water-soluble natively folded protein structure. Unfolding and subsequent protein-protein association in solution is typically accompanied to exposure of hydrophobic core to the solvent. In proteins that associate with membranes, exposure of hydrophobic sites to the solvent is expected to increase a protein membrane-association propensity. In diphtheria toxin increase of the membrane propensity is assumed at low pH due to partial unfolding and refolding of a protein into a postulated membrane competent state. Such a state may be characterized by increased exposure of hydrophobic sites and partial loss of hydrophobic core. In this subsection we analyze how structural changes observed in the simulations and characterized in previous subsections affect solvent exposure of hydrophobic site. From analysis of T1 simulation in the previous sections we identified the non-polar helix TH8 as the T-domain core surrounded by hydrophobic surfaces of helices TH3, TH5 and TH9. The neutral pH MD simulation shows a stable core comprised of helices TH3, TH5 and TH8 (RMSF and RMSD less than 1 Å, see Figure 5 and Figure 8A, respectively). In contrast, C_α_-RMSD traces increase to ca. 1.3 Å and 2.2 Å over both low pH trajectories T2 and T3, respectively. In both trajectories this increase is due to N-terminal unfolding of TH5. This unfolding occurs in the vicinity of protonated H251 (helix TH3), which has a brief increase of its side-chain solvent exposure in both trajectories T2 and T3. In trajectory T3, unfolding of a TH5 helical turn results in structural changes near W281 side-chain, which is contained in TH5 as well as Y278 (see Figure S12). Furthermore, Y278 adopts a new preferred state involving side-chain/side-chain interactions with protonated H251, which is accompanied by disruption of hydrogen bonds (Y278-S332 and Y278-S336). Previous sections have shown that TH9 undergoes partial C-terminal unfolding in both low pH T-domain MD runs. Therefore, to characterize the dynamics of TH9 relative to the stable helix TH8, we determined their inter-helical crossing angle. This is calculated using C_α_ atoms of the stable TH9 N-terminus. Trajectory T3 shows a larger average angle (57 °) than the average angle over trajectory T2 (41 °) and the one obtained over the neutral pH protein trajectory (44 °). The inter-helical angle in trajectory T3 is characterized by an early transition to around 60 °, in contrast to the late transition to larger angles over the last 400 ns of trajectory T2, see Figure 8B. This angle increase coincides with the C_α_-RMSD increase of the hydrophobic core formed by helices TH3, TH5 and TH8, as shown in Figure 8A. Helix axes were determined using C_α_ atoms with RMSF < 1.3 Å in all MD simulations. Similar calculations show that helical packing of TH6 and TH9 is highly dynamic in all MD trajectories. The inter-helical angles vary between 48 ° to 90 ° with an approximate average angle of 72 ° in all MD simulations. This core is maintained with an average C_α_-RMSD < 1.7 Å in all MD simulations. The crossing angle fluctuations of helix TH9 highlights the loosening of hydrophobic contacts and solvent exposure of hydrophobic regions of the putative T-domain trans-membrane alpha-helices in conditions that model low pH solution. The following section describes these changes observed in our T-domain simulations.

**Figure 8.**
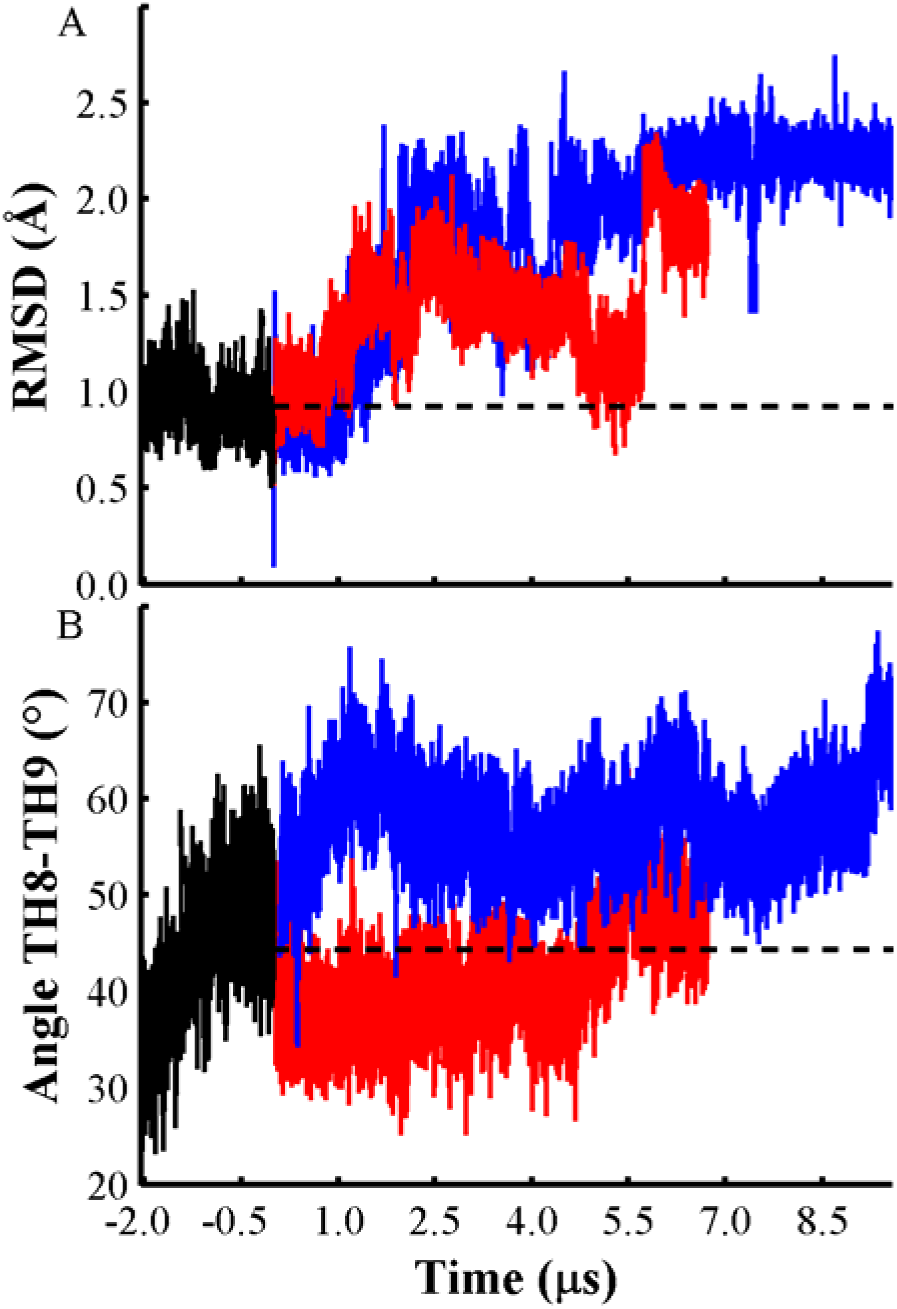
(A) C_α_-RMSD of residues involved in the hydrophobic core formed by TH3, TH5 and TH8. RMSD traces trajectory T1 (black line), T2 (red line), and T3 (blue). (B) Inter-helical angle between helices TH8 and TH9 along trajectories T1 (black), T2 (red), and T3 (blue). Broken black lines represent the average RMSD (0.9 Å) and angle (44 °) calculated from T1.

To identify changes on the protein surface hydrophobicity we calculated the hydrophobic solvent accessible surface area (HP-SASA). There is an average increase of approximately 400 Å^2^ of HP-SASA in both low pH MD trajectories relative to the neutral pH MD simulation (see Figure S13A). In particular, helices TH8 and TH9 have more hydrophobic sites exposed to the solvent in the low pH T-domain simulations than the rest of the protein. The HP-SASA of TH9 has an average value of 116 Å^2^ through the T1 trajectory. This average increases to 187 Å^2^ and 213 Å^2^ over T2 and T3 trajectories, respectively. HP-SASA of TH8 has an average value of 39.3 Å^2^ calculated over the T1 trajectory. In contrast, this average value increases to 97.3 Å^2^ over the T2 trajectory and it remains around 40.7 Å^2^ over the T3 trajectory.

Several hydrophobic sites in helix TH9 that are located in the hydrophobic interface of helices TH8 and TH9 become solvent exposed. For example, the F368 side-chain is significantly buried in trajectory T1 (8.4 Å^2^), but its solvent accessibility increases along T2 (16.6 Å^2^) and T3 (36.6 Å^2^); the V371 side-chain remains partially buried in trajectory T1 (10.2 Å^2^), but it is greatly exposed to the solvent over both low pH T domain trajectories T2 (40.2 Å^2^) and T3 (78.5 Å^2^); and I365 side-chain has a higher average SASA value in trajectory T2 (22.8 Å^2^) than in T1 (11.0 Å^2^) (yet, is it almost buried in T3 (1.3 Å^2^).

In addition to the increased solvent exposure of the hydrophobic regions in helices TH8 and TH9 in T2 and T3 other protein regions become more solvent accessible. The SASA of the entire protein remains 9161 Å^2^ on average over the neutral pH T-domain simulation (T1) and increases to 9639 Å^2^ and 9552 Å^2^ in the trajectories T2 and T3, respectively (see Figure S13B). Analysis of the different contributions to the protein solvent exposure shows that refolding of helices TH1 and TH2, as well as loop fluctuations contribute to the increased SASA in both low pH T-domain MD simulations.

## 4. CONCLUSIONS

Partially unfolded ensembles of a diphtheria toxin T-domain model at low pH conditions were generated in two independent multi-microsecond MD simulations. The third simulation presented here is of a protein in a folded form at neutral pH. In both low pH simulations protein unfolding transitions occurred in the time-frame between 1 µs to 5 µs. Further simulations led to formation of partially refolded structures with overall compact structure. These partially unfolded (from a native structure) and refolded structures may be attributed to formation of a membrane competent state of the protein, which has been identified in previous work based on experiments of Ladokhin and co-workers.^19,21^

The neutral pH T-domain simulation shows that the T-domain structure is stable, but it exhibits the unfolding and refolding of several helices. It also shows two different neutral pH T-domain states, which are characterized by the disruption of a salt-bridge in helices TH1 and TH9. Both low pH T-domain simulations show partial unfolding of the protein within 2 µs of the simulations. Both T2 and T3 refolded structures are stabilized over the last microseconds of the simulations in compact conformations with low RMSFs. The simulated meta-stable state of a protein shows increased protein core fluctuations, increased solvent accessible surface of the protein, as well as increased solvent exposure of hydrophobic regions, which may facilitate protein-membrane interactions through the T-domain membrane insertion pathway.

PCA analysis was applied to various metrics that characterize conformational structural ensembles. The similarities of the conformational space sampled in our independent trajectories were best resolved using the helicity measure, earlier introduced by Speransky et al.,^33^ In this measure, the helicity along the primary sequence is determined using three consecutive residues. The helicity measure shows that trajectories T2 and T3 sample similar conformational subspace in the space described by the two lowest principal components of the joint dataset (T1, T2). In these principal components the structures show partial subspace overlap of the last stable segments of T2 and T3 trajectories. Analysis of individual contribution of each residue triplet to the slowest principal component PC1 shows that in both unfolding trajectories T2 and T3 the unfolding transition and the resulting re-folded conformation is mainly influenced by triplets containing H257. In general, the observed importance of H257 is in good agreement with its suggested role in destabilization of T-domain structure at low pH.^21,24^ The second most important principal component, PC2 is influenced by the residues in helices TH6 or TH7 and by those near H223, which is located in the partially refolded N-terminal region. Thus, PCA analysis using the helicity measure based on triplets of residues is useful in the description of the dynamics of partially unfolded proteins. In contrast, the PCA analysis on the backbone dihedral space (a metric that is often used in characterization of protein structural ensembles) is unable to capture relative importance of different amino acids in the slow conformational changes observed in T-domain. Overall, the PCA analysis of trajectories T2 and T3 describes two similar transition routes from folded to partially unfolded T-domain ensembles.

Our simulations indicate that protonation of histidines triggers the refolding/unfolding of N-terminal helices TH1 and TH2. Local conformational changes in helix TH1 were also detected in titration experiments by Wang et al.,^22^ in which single fluorescence labels spanning helix TH1 suggested solvent exposure of hydrophobic residues in TH1. To rotate this helix, which was proposed by the authors, it may be necessary to break inter-helical salt bridges. As a result of histidines protonation, several inter-helical salt-bridges located in the protein N-terminal are disrupted in our simulations of low pH T-domain in solution. For example, the salt bridge involving residue K216, which indeed occurs in both trajectories T2 and T3. Conformational flip of the K216 backbone dihedral angle initiates unfolding of TH1. In presence of lipid membranes disruption of these salt-bridges may facilitate protein-membrane interactions.

Helix TH2 unfolds rapidly in both low pH protein simulations. This helix is predicted to be disordered by a consensus of disorder prediction algorithms.^44^ Thus, our molecular dynamics studies coupled with disorder predictors show the role of protonation in triggering disordered regions of otherwise structurally stable proteins at neutral pH. The protonation-dependent disorder of TH2 may have implications for the membrane association and subsequent translocation of the protein N-terminus across the lipid bilayer. A similar case of locally disordered protein triggered by protonation of a single residue has been recently observed in multiple microsecond long dynamics of EGFR.^31^

Unfolding transition of helix TH1 follows conformational changes of TH4 in both low pH simulations. This can be explained by the loss of multiple tertiary contacts of these helices and helix TH8 and the disruption of a hydrogen bond network between residues in TH1 and H257 in the inter-helical loop of TH3-4. Previously, we have reported a significant free energy change of protonation of H257 in the protein native state (6.9 kcal/mol),^24^ which may explain significant backbone rearrangements required to accommodate a positive charge in H257 side-chain. The positively charged side-chains of H223 and H257 repeatedly increase their distance between themselves in both low pH trajectories. The interaction between this pair of histidines has been suggested as a major feature of the pH-dependent function of T-domain^24^ and other molecular systems in nature.^45^ Similarly, a positive free energy of protonation of H251 in the native state (3.7 kcal/mol),^24^ may explain the observed conformational changes in helix TH5 N-terminus, which is near the positively charged H251. A partially unfolded full toxin crystal structure^23^ shows helix TH4 unfolded, partial unfolding of TH1 and TH5 and the lack of electron density of TH2, which are in agreement with our simulations. Also, partial loss of helicity of TH1 at K216 is in agreement with the simulations.

In summary, we have presented extensive MD simulations and detailed analysis of partial unfolding of the diphtheria toxin T-domain upon protonation of histidine residues in solution. Thorough comparison of simulated structures with experimental data indicates that the results of the simulations complement experimental investigation to provide unattainable microscopic details of structures that undergo transient modifications. Molecular modeling thus proves to be a valuable tool for such systems. There are practical limitations in any MD simulation that may adversely affect simulated structures. For example, small simulation box sizes were indicated to affect the protein dynamics in explicit solvent simulations.^46,47^ In the present work, to assess the severity of this limitation for the predicted protein unfolding, we designed simulations with two different box sizes. Trajectory T3 is a simulation with a larger simulation box exhibited very similar unfolded ensemble comparing to T2 (a smaller box simulation), especially at similar times. Further modification of the structure in T3 simulation was observed due to its longer simulation time. Moreover, at intermediate time, the structures in T3 simulation were more compact than in T2 simulation, indicating that the box size did not preclude the protein from unfolding and reconfiguring its conformation.

Here we used a pre-determined fixed protonation state of a protein to mimic its state at a specific pH. This is an approximation which allows us to simulate protein behavior in real time over microsecond long time-scales. A potentially better approach would be to perform a constant pH molecular dynamics, in which protonation states of protonatable residues can be modified over simulation time according to their current pK_a_ values. However, such approach is currently unattainable for the simulation times needed to unfold a protein and to achieve equilibrated protonation states of all residues. Development of biased molecular dynamics simulations may as well enhance generation of ensembles of partially unfolded proteins in solution, and this is a subject of future work.

## ASSOCIATED CONTENT

### Supporting Information

Contact maps, overlay of helices TH1-2 from representative structures, K216 backbone dihedral angle trace versus time, histograms of inter-helical salt-bridges disrupted, principal component analysis on backbone dihedral space, principal components variance contribution in triplet and dihedral space, influence of triplets and dihedral angles on the two first principal components, conformational change of side-chain Y278. This material is available free of charge via the Internet at http://pubs.acs.org.

## AUTHOR INFORMATION

### Notes

The authors declare no competing financial interest.

## ACKNOWLEDGMENT

We are grateful to Dr. A.S. Ladokhin for helpful discussions. This research was supported by National Institutes of Health grant GM-069783. Computational work was in part done on ANTON supercomputer supported under NIH RC2GM093307 grant, as well as TGMCB040051N XSEDE grant.

## REFERENCES

(1) Collier, R. J. Toxicon 2001, 39, 1793.

(2) Murphy, J. R. Toxins 2011, 3, 294.

(3) Ladokhin, A. S. Toxins 2013, 5, 1362.

(4) Honjo, T.; Nishizuk. Y; Hayaishi, O.; Kato, I. Journal of Biological Chemistry 1968, 243, 3553.

(5) Shiver, J. W.; Donovan, J. J. Biochimica Et Biophysica Acta 1987, 903, 48.

(6) Oh, K. J.; Zhan, H. J.; Cui, C.; Hideg, K.; Collier, R. J.; Hubbell, W. L. Science 1996, 273, 810.

(7) Zimmer, J.; Nam, Y. S.; Rapoport, T. A. Nature 2008, 455, 936.

(8) Koriazova, L. K.; Montal, M. Nature Structural Biology 2003, 10, 13.

(9) Prince, H. M.; Duvic, M.; Martin, A.; Sterry, W.; Assaf, C.; Sun, Y. J.; Straus, D.; Acosta, M.; Negro-Vilar, A. Journal of Clinical Oncology 2010, 28, 1870.

(10) Bennett, M. J.; Choe, S.; Eisenberg, D. Protein Science 1994, 3, 1444.

(11) Steere, B. Characterization of high-order oligomerization and energetics in Diptheria Toxin, University of California, 2001.

(12) Senzel, L.; Gordon, M.; Blaustein, R. O.; Oh, K. J.; Collier, R. J.; Finkelstein, A. Journal of General Physiology 2000, 115, 421.

(13) Rosconi, M. P.; Zhao, G.; London, E. Biochemistry 2004, 43, 9127.

(14) Zhao, G.; London, E. Biochemistry 2005, 44, 4488.

(15) Chenal, A.; Prongidi-Fix, L.; Perier, A.; Aisenbrey, C.; Vernier, G.; Lambotte, S.; Haertlein, M.; Dauvergne, M. T.; Fragneto, G.; Bechinger, B.; Gillet, D.; Forge, V.; Ferrand, M. J Mol Biol 2009, 391, 872.

(16) Murphy, R. F.; Powers, S.; Cantor, C. R. Journal of Cell Biology 1984, 98, 1757.

(17) Chenal, A.; Savarin, P.; Nizard, P.; Guillain, F.; Gillet, D.; Forge, V. J Biol Chem 2002, 277, 43425.

(18) Ladokhin, A. S.; Legmann, R.; Collier, R. J.; White, S. H. Biochemistry 2004, 43, 7451.

(19) Kyrychenko, A.; Posokhov, Y. O.; Rodnin, M. V.; Ladokhin, A. S. Biochemistry 2009, 48, 7584.

(20) Perier, A.; Chassaing, A.; Raffestin, S.; Pichard, S.; Masella, M.; Menez, A.; Forge, V.; Chenal, A.; Gillet, D. Journal of Biological Chemistry 2007, 282, 24239.

(21) Rodnin, M. V.; Kyrychenko, A.; Kienker, P.; Sharma, O.; Posokhov, Y. O.; Collier, R. J.; Finkelstein, A.; Ladokhin, A. S. Journal of Molecular Biology 2010, 402, 1.

(22) Wang, J.; Rosconi, M. P.; London, E. Biochemistry 2006, 45, 8124.

(23) Leka, O.; Vallese, F.; Pirazzini, M.; Berto, P.; Montecucco, C.; Zanotti, G. FEBS J 2014.

(24) Kurnikov, I. V.; Kyrychenko, A.; Flores-Canales, J. C.; Rodnin, M. V.; Simakov, N.; Vargas-Uribe, M.; Posokhov, Y. O.; Kurnikova, M.; Ladokhin, A. S. J Mol Biol 2013, 425, 2752.

(25) Simonson, T.; Carlsson, J.; Case, D. A. J Am Chem Soc 2004, 126, 4167.

(26) Shaw, D. E.; Deneroff, M. M.; Dror, R. O.; Kuskin, J. S.; Larson, R. H.; Salmon, J. K.; Young, C.; Batson, B.; Bowers, K. J.; Chao, J. C.; Eastwood, M. P.; Gagliardo, J.; Grossman, J. P.; Ho, C. R.; Ierardi, D. J.; Kolossvary, I.; Klepeis, J. L.; Layman, T.; Mcleavey, C.; Moraes, M. A.; Mueller, R.; Priest, E. C.; Shan, Y. B.; Spengler, J.; Theobald, M.; Towles, B.; Wang, S. C. Communications of the Acm 2008, 51, 91.

(27) Shaw, D. E.; Maragakis, P.; Lindorff-Larsen, K.; Piana, S.; Dror, R. O.; Eastwood, M. P.; Bank, J. A.; Jumper, J. M.; Salmon, J. K.; Shan, Y.; Wriggers, W. Science 2010, 330, 341.

(28) Hornak, V.; Abel, R.; Okur, A.; Strockbine, B.; Roitberg, A.; Simmerling, C. Proteins 2006, 65, 712.

(29) Lindorff-Larsen, K.; Maragakis, P.; Piana, S.; Eastwood, M. P.; Dror, R. O.; Shaw, D. E. PLoS One 2012, 7, e32131.

(30) Lindorff-Larsen, K.; Trbovic, N.; Maragakis, P.; Piana, S.; Shaw, D. E. J Am Chem Soc 2012, 134, 3787.

(31) Shan, Y.; Eastwood, M. P.; Zhang, X.; Kim, E. T.; Arkhipov, A.; Dror, R. O.; Jumper, J.; Kuriyan, J.; Shaw, D. E. Cell 2012, 149, 860.

(32) Shan, Y.; Arkhipov, A.; Kim, E. T.; Pan, A. C.; Shaw, D. E. Proc Natl Acad Sci U S A 2013, 110, 7270.

(33) Speranskiy, K.; Kurnikova, M. G. Proteins-Structure Function and Bioinformatics 2009, 76, 271.

(34) Case, D. A.; Darden, T. A.; Cheatham, T. E., III; Simmerling, C. L.; Wang, J.; Duke, R. E.; Luo, R.; Merz, K. M.; Pearlman, D. A.; Crowley, M.; Walker, R. C.; Zhang, W.; Wang, B.; Hayik, S.; Roitberg, A.; Seabra, G.; Wong, K. F.; Paesani, F.; Wu, X.; Brozell, S.; Tsui, V.; Gohlke, H.; Yang, L.; Tan, C.; Mongan, J.; Hornak, V.; Cui, G.; Beroza, P.; Mathews, D. H.; Schafmeister, C.; Ross, W. S.; Kollman, P. A. AMBER 9, University of California, San Francisco: SanFrancisco, CA, 2006.

(35) Ryckaert, J. P.; Ciccotti, G.; Berendsen, H. J. C. J Comput Phys 1977, 23, 327.

(36) Darden, T.; York, D.; Pedersen, L. Journal of Chemical Physics 1993, 98, 10089.

(37) Martyna, G. J.; Tobias, D. J.; Klein, M. L. Journal of Chemical Physics 1994, 101, 4177.

(38) Shan, Y. B.; Klepeis, J. L.; Eastwood, M. P.; Dror, R. O.; Shaw, D. E. Journal of Chemical Physics 2005, 122.

(39) Humphrey, W.; Dalke, A.; Schulten, K. J Mol Graph 1996, 14, 33.

(40) Kabsch, W.; Sander, C. Biopolymers 1983, 22, 2577.

(41) Lange, O. F.; Grubmuller, H. Proteins 2008, 70, 1294.

(42) Amadei, A.; Linssen, A. B.; Berendsen, H. J. Proteins 1993, 17, 412.

(43) Altis, A.; Nguyen, P. H.; Hegger, R.; Stock, G. J Chem Phys 2007, 126, 244111.

(44) Huang, Y. J.; Acton, T. B.; Montelione, G. T. Methods Mol Biol 2014, 1091, 3.

(45) Harrison, J. S.; Higgins, C. D.; O’Meara, M. J.; Koellhoffer, J. F.; Kuhlman, B. A.; Lai, J. R. Structure 2013, 21, 1085.

(46) Takemura, K.; Kitao, A. J Phys Chem B 2007, 111, 11870.

(47) Wassenaar, T. A.; Mark, A. E. J Comput Chem 2006, 27, 316.

