## Supporting Information for "Microsecond Molecular Dynamics Simulations of Diphtheria Toxin Translocation T-Domain pH-Dependent Unfolding in Solution"

**RESULTS**

**Principal Component Analysis on Dihedral Space.**

In addition to the principal component analysis (PCA) on the helical triplet space, we performed PCA analysis on the backbone dihedral angles of the ten α-helices identified in the crystal structure (TH1-9, TH5’). We defined two datasets containing the last 1 µs segments of trajectories (T1, T2) and (T1, T3). Thus, these datasets contain an ensemble of folded and refolded conformations of T-domain. We first calculated the principal components (PC) from the dataset (T1, T2). Figure S8A shows that the last 1 µs segments of T2 and T3 have no overlap in a reduced conformational subspace described by the two lowest principal components. Figure S8C shows the projection of all MD frames. T2 contains the majority of T3 MD frames on the two principal components of dataset (T1, T2), which is similar to our observations in the triplet space (helicity), see Figure 7C. Secondly, we also performed PCA analysis on the dataset (T1, T3). Figure S8B shows no overlap of last segments of T2 and T3 on the first two principal components. Figure S8D shows partial overlapping of the entire trajectories T2 and T3.

The first principal component of both datasets (T1, T2) and (T1, T3) has a significant variance contribution ca. 54 – 56 %, see Figure S9. This contribution of PC1 is similar to that observed in the triplet space. However, residues with the largest influence in PC1 are different for the dihedral and triple space. For example, Fig S10A shows that a backbone dihedral in residue T267 has the largest influence in PC1 for the dataset (T1, T2), while a triplet containing residue H257 in PC1 obtained from the same dataset (see Figure S7A). In general this is also observed for dataset (T1, T3). For example, Figure S10B shows that a backbone dihedral in E298 has the largest influence in PC1, while a triplet containing residue H257 has the largest influence for the same dataset (see Figure S7B).

We also performed dihedral PCA analysis over the complete trajectories T2 and T3. We conclude that the projection of T2 and T3 have no significant overlap on the lowest PC vectors from T2 or T3 (see Figures S11A, B). Notice that PC1 has a smaller variance contribution ca. 18 – 22 % (see Figure S12), which is lower than the ones calculated for datasets (T1, T2) and (T1, T3).

**FIGURES**

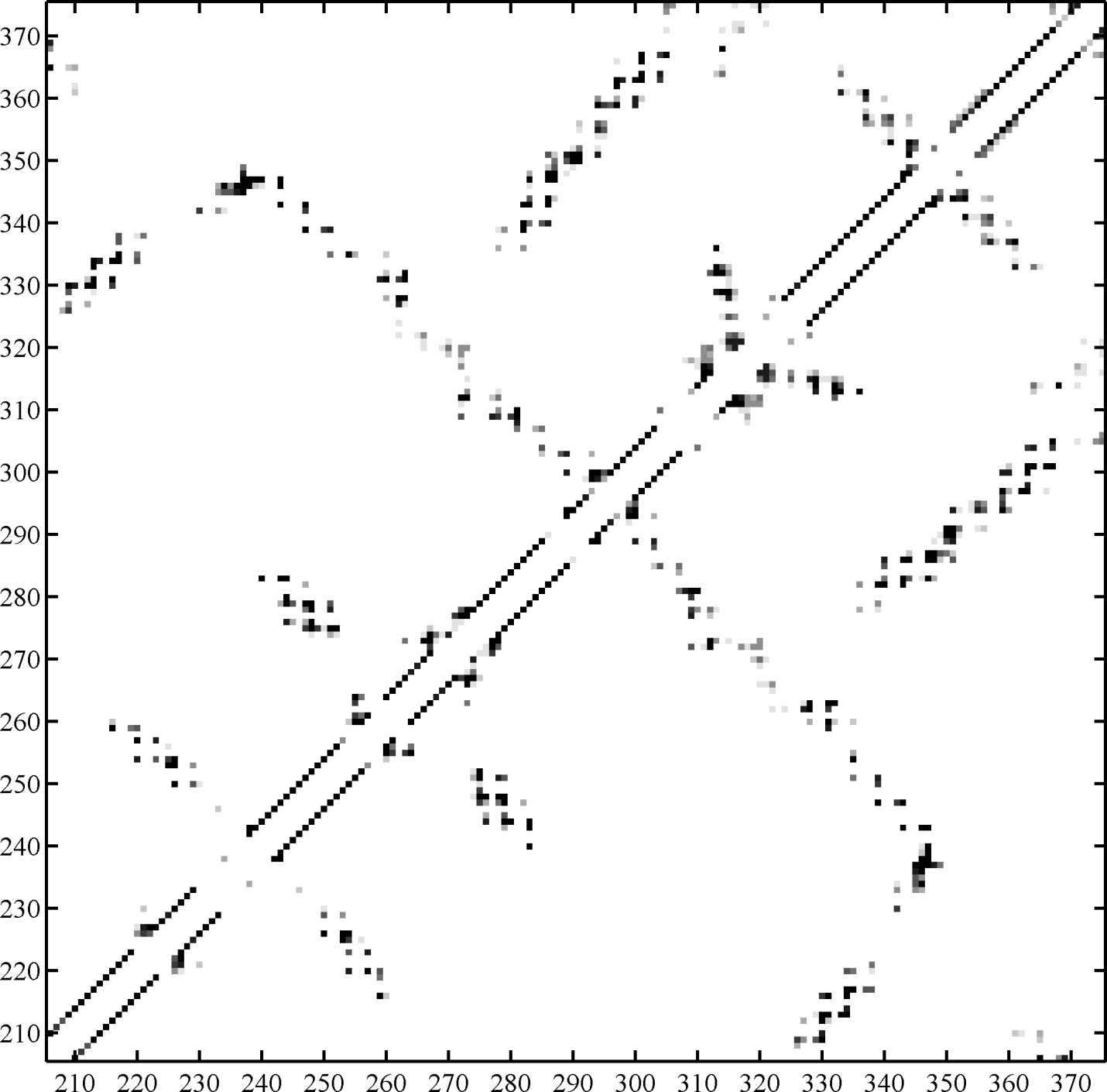

**Figure S1A.** Change of tertiary contacts depicted by the fraction of non-local contacts, calculated over Cα atoms of residues 206-375 in trajectory T1. Fraction values in the range of 0 and 1 are represented by grey scale. Fraction of non-local contacts between pairs ij of Cα atoms separated by at least 8 Å and residues containing atoms i and j are separated by at least 3 residues.

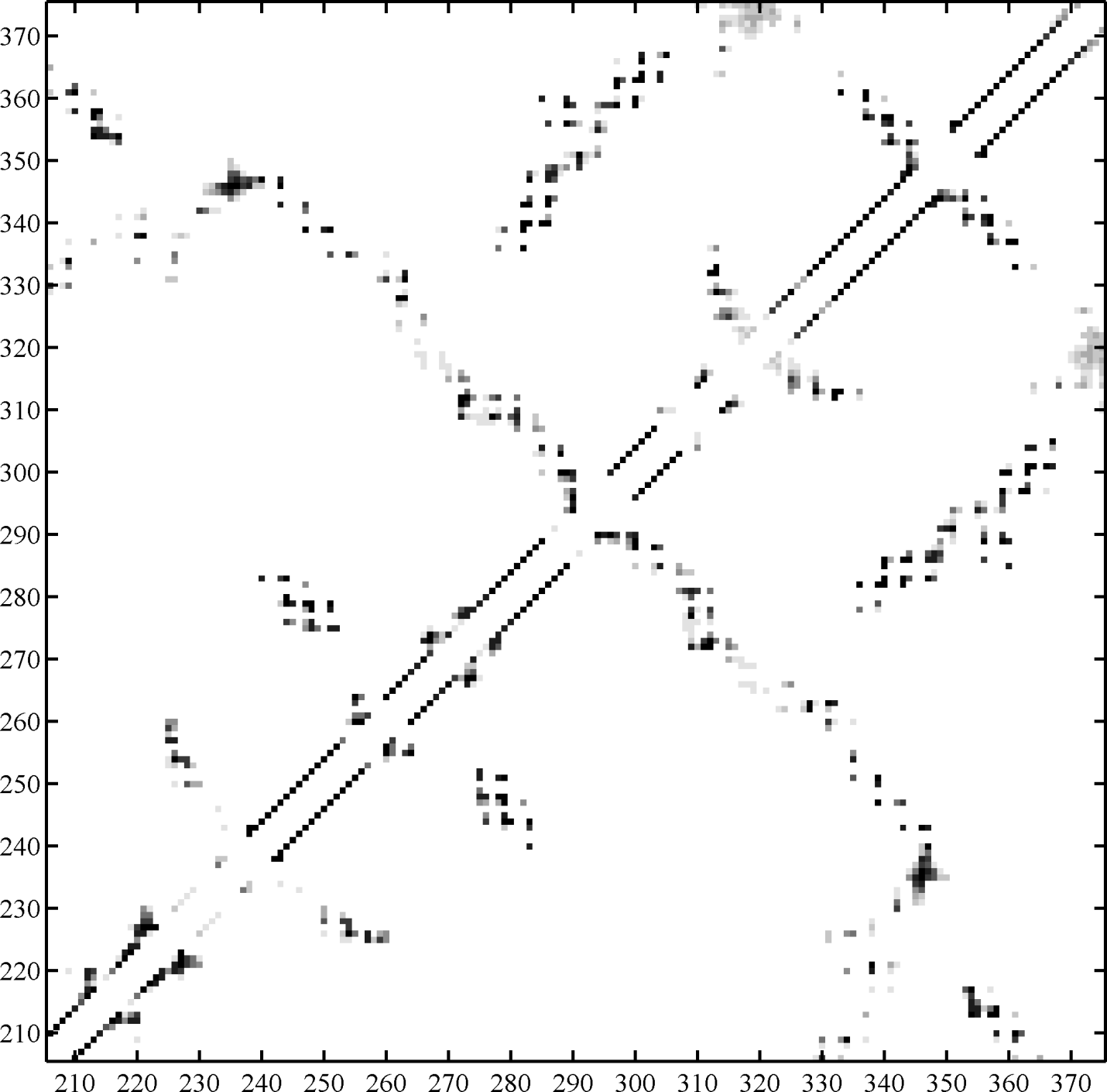

**Figure S1B.** Change of tertiary contacts depicted by the fraction of non-local contacts, calculated over Cα atoms of residues 206-375 in trajectory T2. Fraction values in the range of 0 and 1 are represented by grey scale. Fraction of non-local contacts between pairs ij of Cα atoms separated by at least 8 Å and residues containing atoms i and j are separated by at least 3 residues.

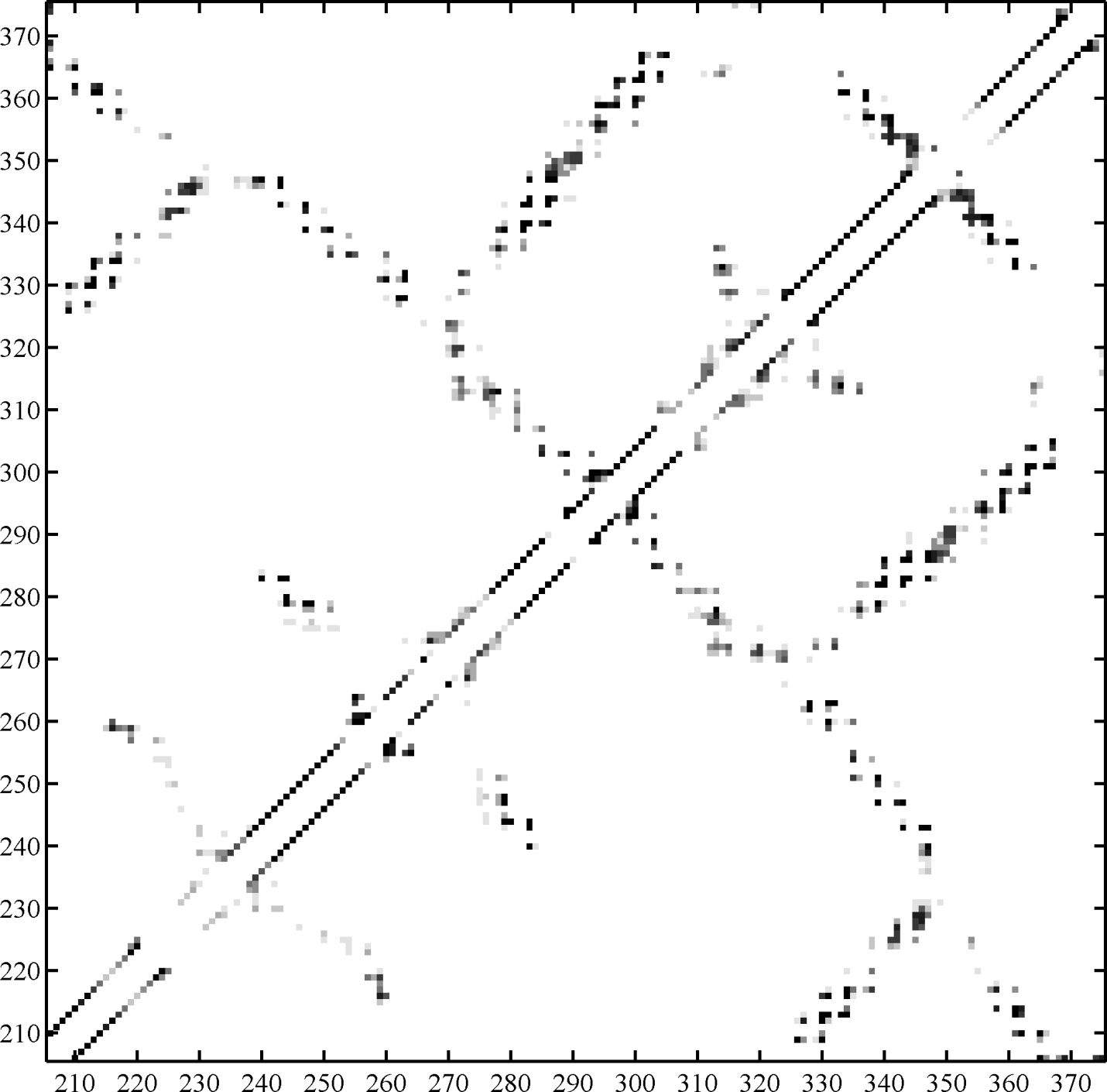

**Figure S1C.** Change of tertiary contacts depicted by the fraction of non-local contacts, calculated over Cα atoms of residues 206-375 in trajectory T3. Fraction values in the range of 0 and 1 are represented by grey scale. Fraction of non-local contacts between pairs ij of Cα atoms separated by at least 8 Å and residues containing atoms i and j are separated by at least 3 residues.

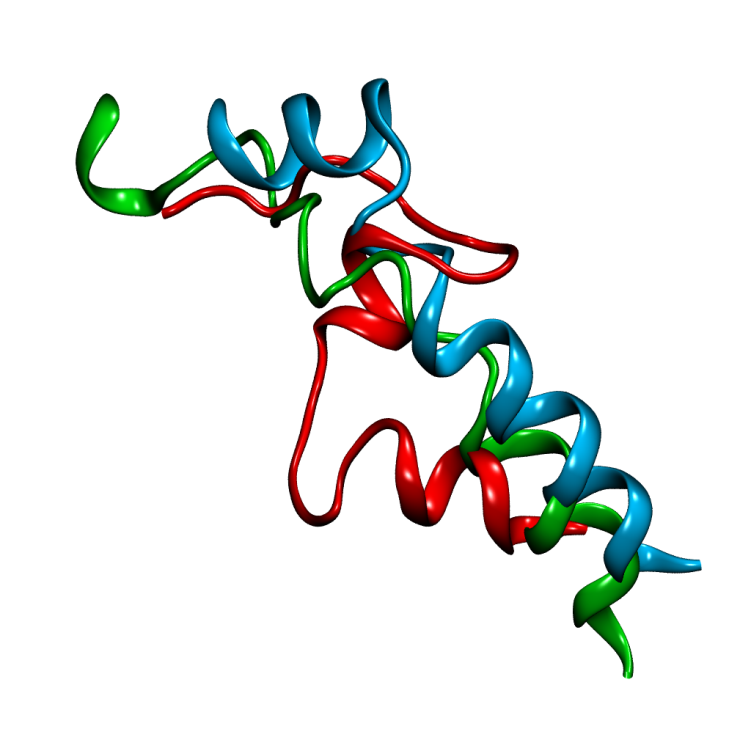

**Figure S2.** Overlay of helices TH1-2 from representative structures obtained from the last 1 µs of trajectories T1 (cyan), T2 (red) and the last 2 µs of T3 (green) in cartoon representation. Helices TH1 and TH2 are partially unfolded in both representatives T2 and T3 after the conformational changes described in the results section.

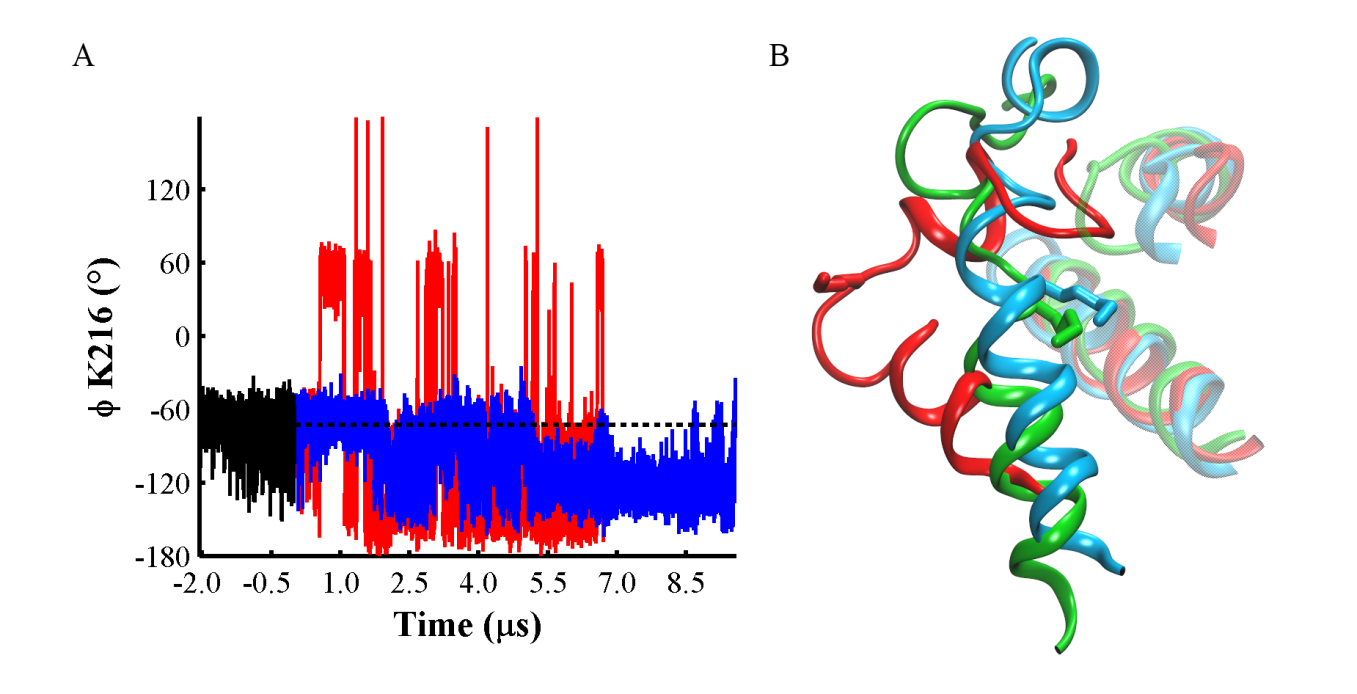

**Figure S3.** (A) φ dihedral angle traces of K216 over trajectories T1 (black line), T2 (red) and T3 (blue). Broken black lines represent the average value of φ K216 (-73 °) in trajectory T1. (B) Overlay of helix TH1 from representative structures of T1 (cyan ribbon), T2 (red) and T3 (green). K216 is highlighted by stick representation. Helices TH3 and TH8 are shown in transparent representation using the same colors.

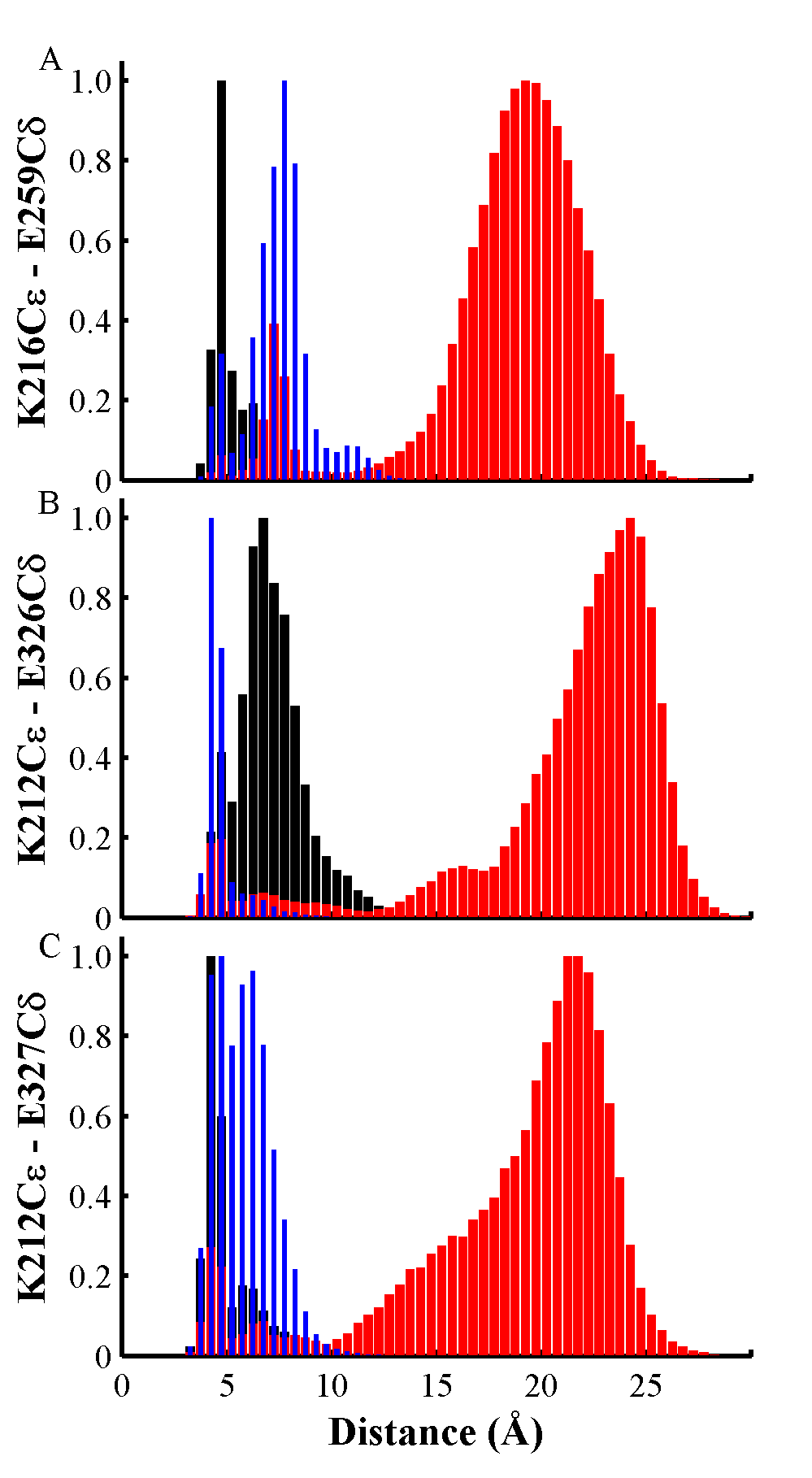

**Figure S4.** Normalized frequency of distances in salt-bridges obtained over trajectories T1 (black histograms), T2 (red histograms), and T3 (blue histograms). (A) Histogram of distances of atoms in side-chains of K216 – E259. (B) Histogram of distances of atoms in side-chains of K212 – E326. C. Histogram of distances of atoms in side-chains of K212 – E327. K212 is located in helix TH1, E259 is located in the loop between TH3 and TH4, E326 and E327 are located in helix TH8.

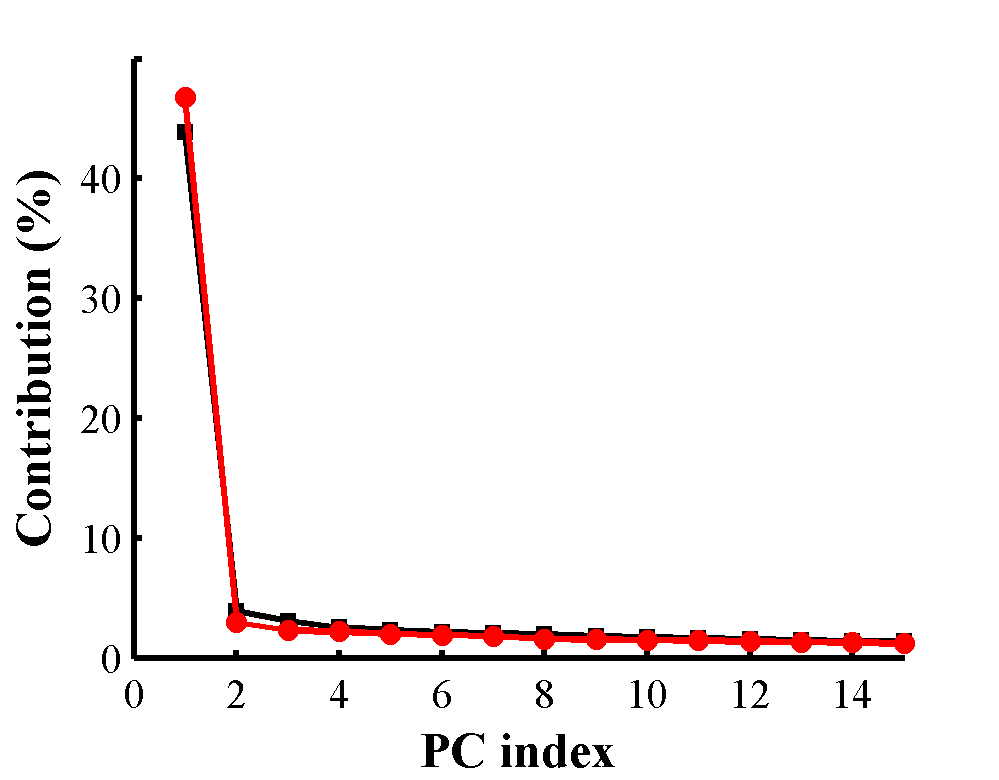

**Figure S5.** Variance contribution of the first 15 principal components (PC) obtained from datasets containing the last 1 µs segments of trajectories (T1, T2) or (T1, T3) shown by black and red lines, respectively. Helicity measure is calculated for each MD frame (residues 206-375).

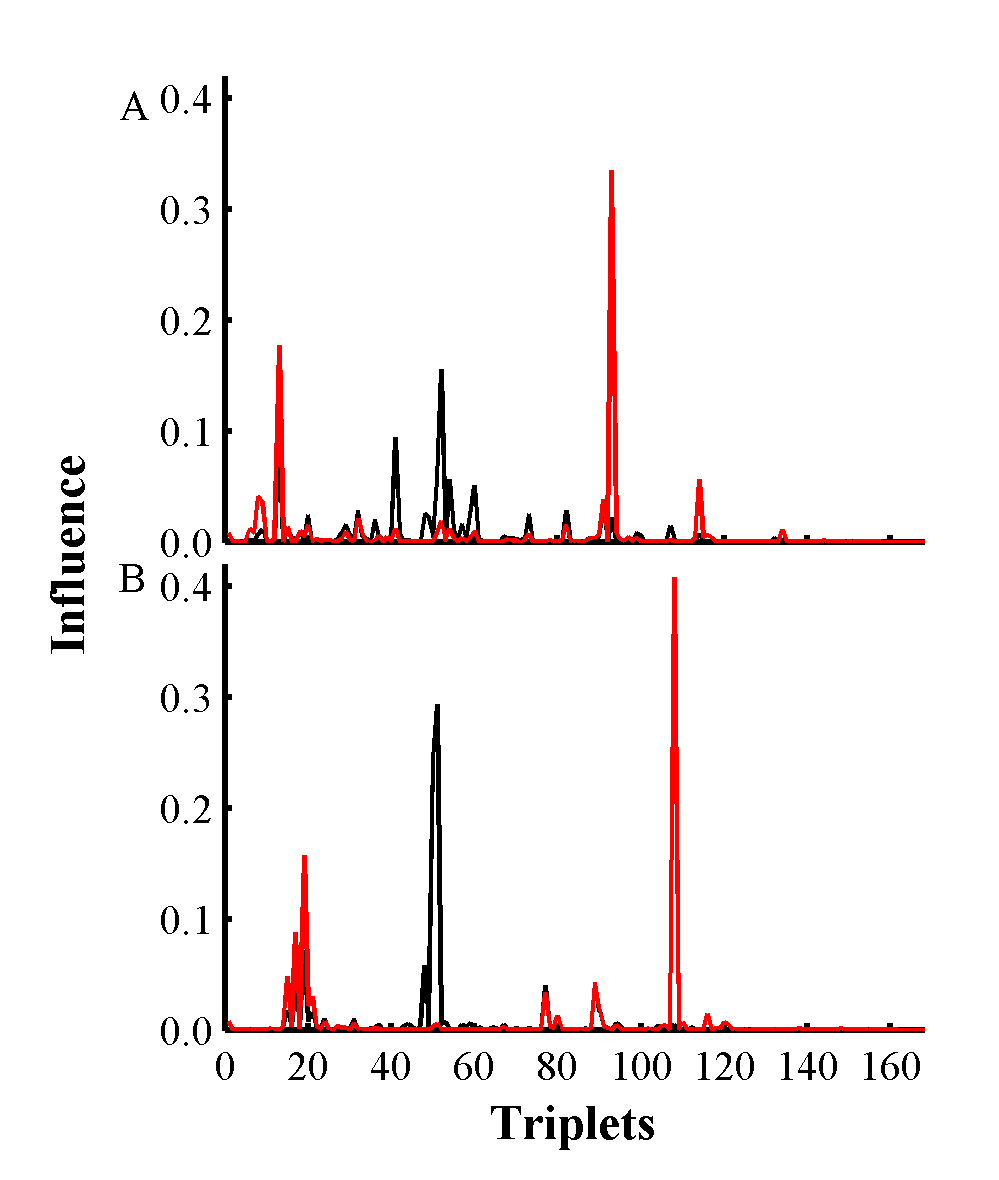

**Figure S6.** Influence of each triplet on the first principal component (black lines) and second principal component (red lines). (A) Influence of each triplet obtained from datasets containing the last 1 µs segments of trajectories (T1, T2). (B) Influence of each triplet obtained from datasets containing the last 1 µs segments of (T1, T3). Helicity measure is calculated for each MD frame (residues 206-375).

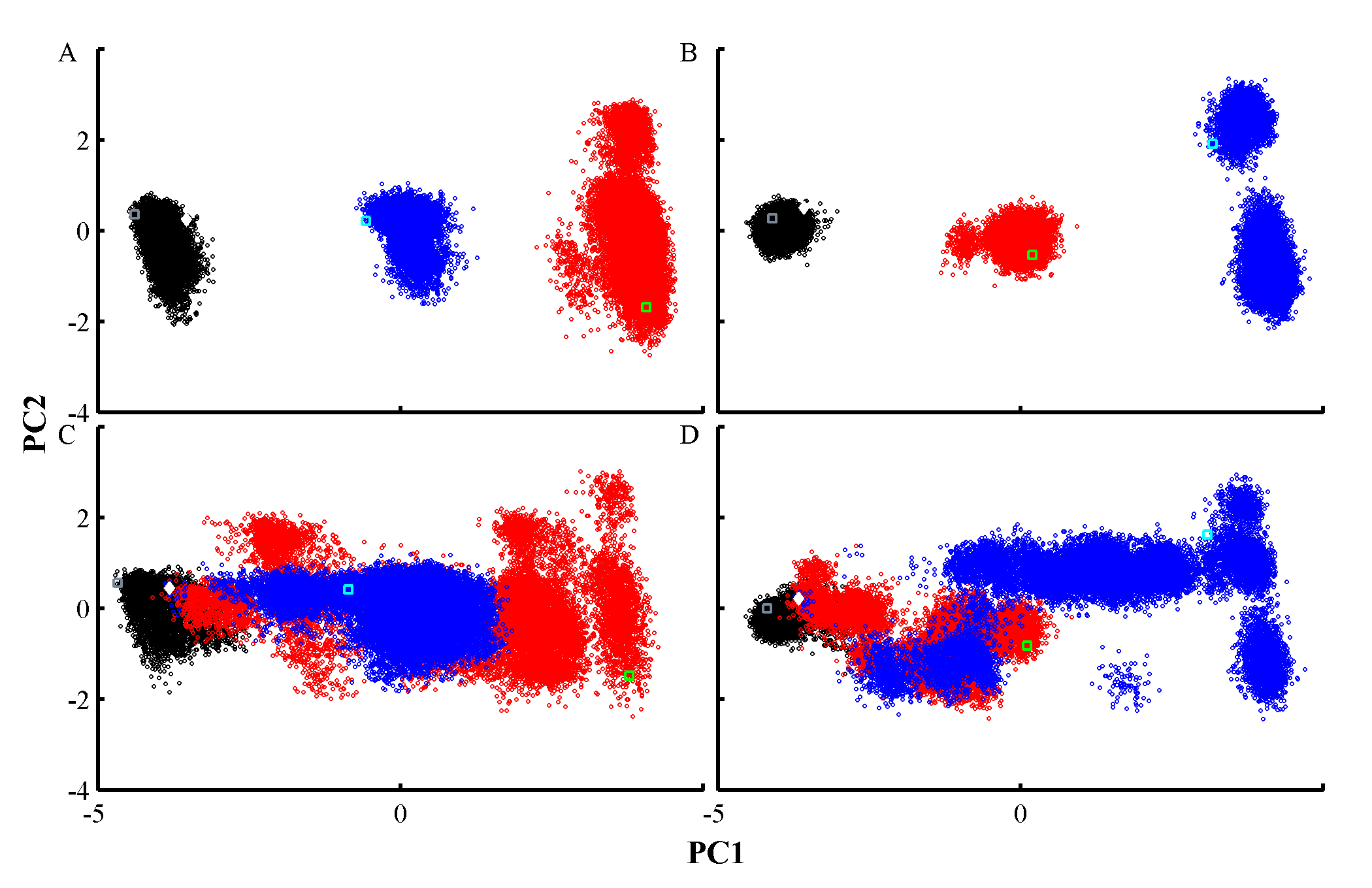

**Figure S7.** Projection of MD trajectories on the two main principal components calculated for two datasets comprising the last 1 µs segments of trajectories (T1, T2) or (T1, T3). MD frames from trajectories T1, T2, and T3 are represented by black, red, and blue circles, respectively. (A) Two-dimensional projection of the last 1 µs of all trajectories on principal components obtained from a dataset containing the last 1 µs of (T1, T2). (B) Two-dimensional projection of the last 1 µs of all trajectories on principal components obtained from a dataset containing the last 1 µs of (T1, T3). (C) Two-dimensional projection of all trajectories on principal components obtained from a dataset containing the last 1 µs of (T1, T2). (D) Two-dimensional projection of all trajectories on principal components obtained from a dataset containing the last 1 µs of (T1, T3). Projection of the X-ray structure is shown in filled white diamond. Final MD frames of trajectories T1, T2, and T3 are shown in grey, cyan, and green empty rectangles, respectively. Dihedral principal component analysis (dPCA) was carried out over the backbone dihedral angles of helices TH1-9, TH5’ identified in the crystal structure, excluding loops and terminal residues.

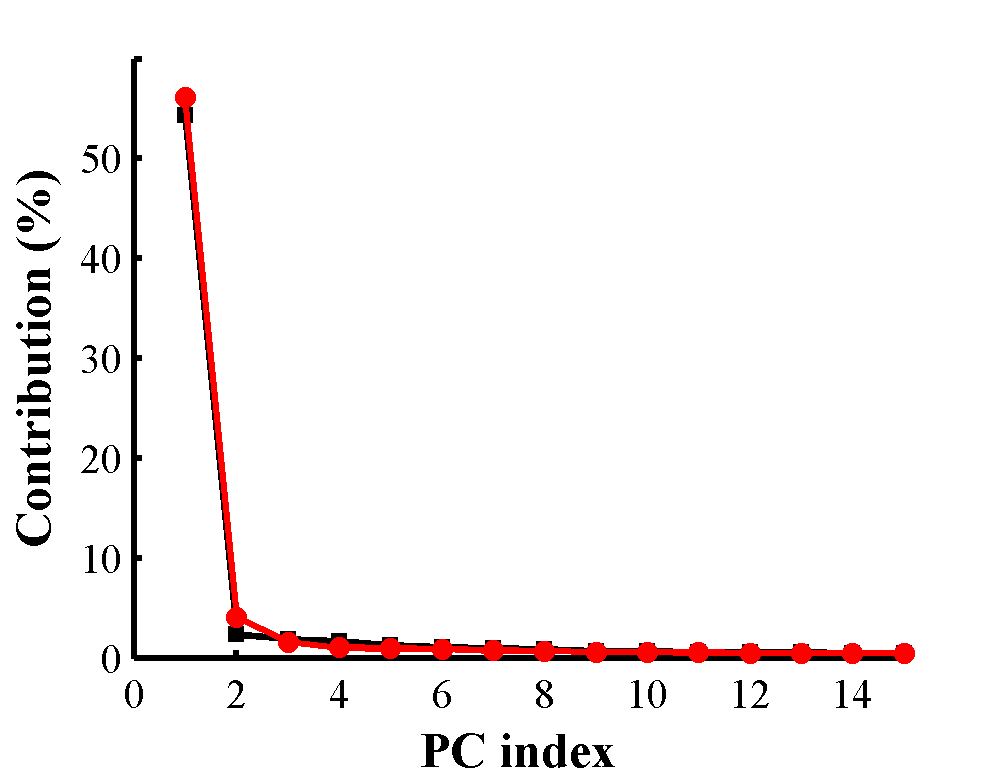

**Figure S8.** Variance contribution of the first 15 principal components (PC) obtained from datasets containing the last 1 µs segments of trajectories (T1, T2) or (T1, T3), shown in black and red lines, respectively. Dihedral PCA analysis was carried out over backbone dihedral angles of residues from helices (TH1-9 and TH5’).

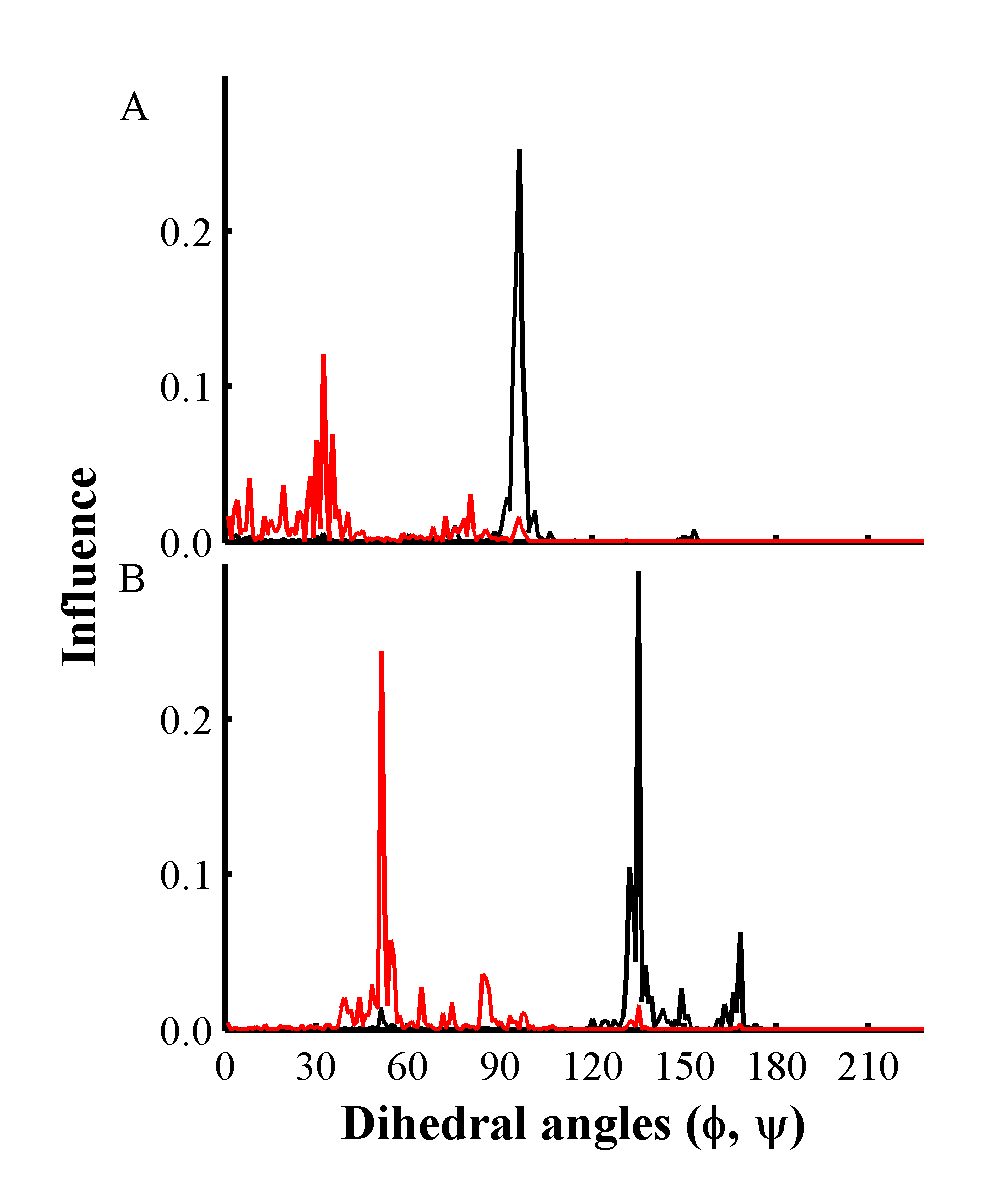

**Figure S9.** Influence of each backbone dihedral angle on the first principal component (black lines) and second principal component (red line). (A) Influence of dihedral angles (φ, ψ) obtained from datasets containing the last 1 µs segments of trajectories (T1, T2). (B) Influence of dihedral angles (φ, ψ) obtained from datasets containing the last 1 µs segments of (T1, T3). Backbone dihedral angles are obtained from residues from helices (TH1-9 and TH5’).

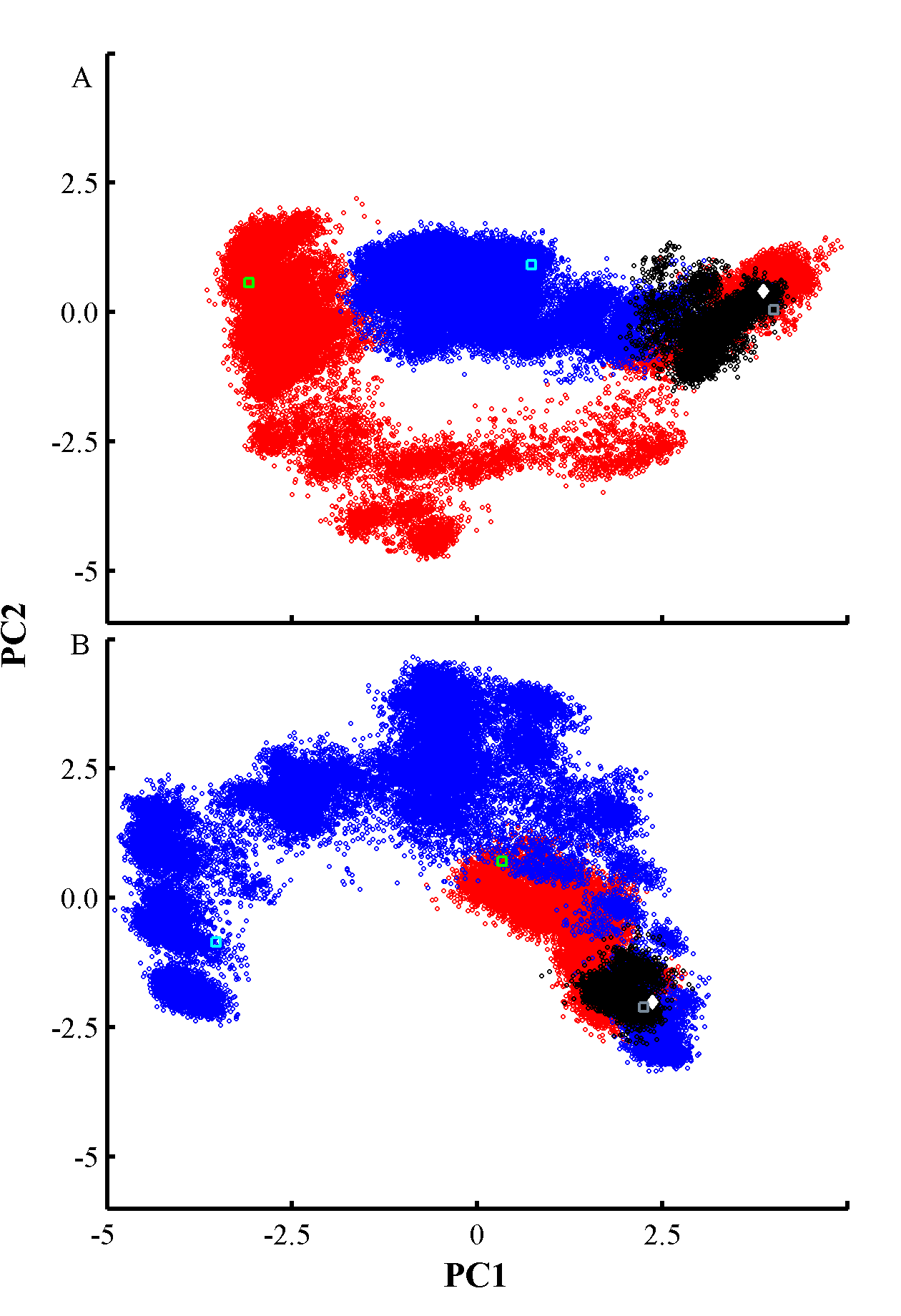

**Figure S10.** Projection of all MD trajectories on the two first principal components calculated using MD frames from: (A) Trajectory T2. (B) Trajectory T3. Dihedral PCA analysis was carried out over backbone dihedral angles of residues from helices (TH1-9 and TH5’), excluding loops and terminal residues. MD snapshots from trajectory T1, T2, and T3 are represented by black, red, and blue circles, respectively. Projection of the X-ray structure is shown in filled white diamond. Final conformations of trajectories T1, T2, and T3 are shown in grey, cyan, and green empty rectangles, respectively.

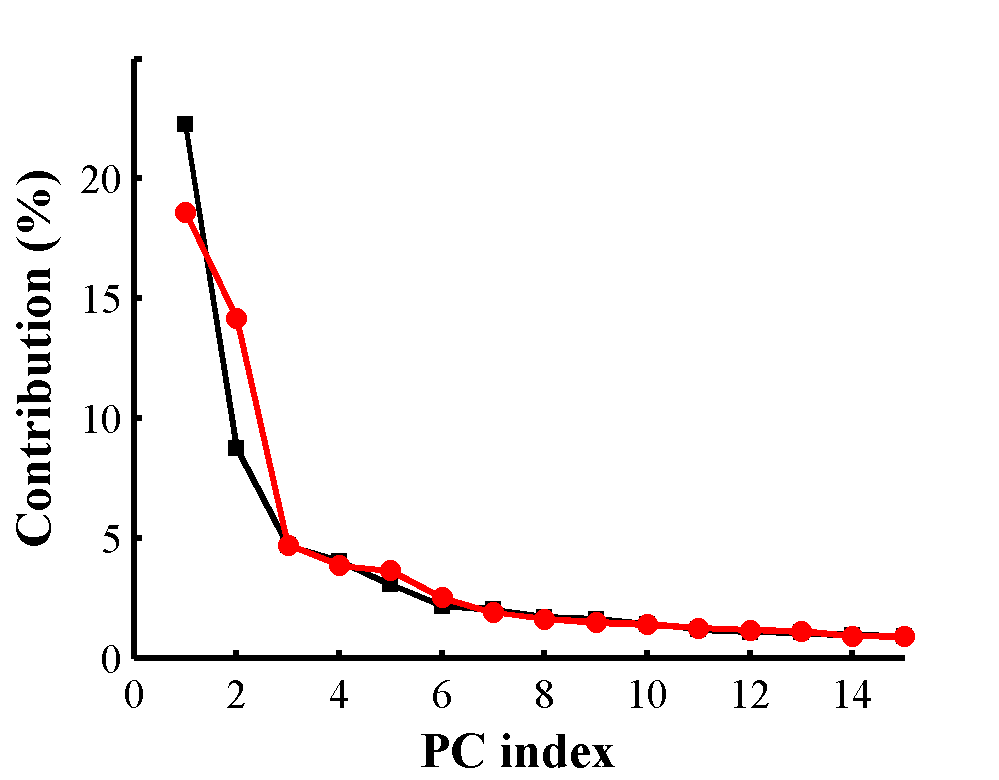

**Figure S11.** Variance contribution of the first 15 principal components obtained from trajectories T2 (black line) and T3 (red line). Dihedral PCA analysis was carried out over backbone dihedral angles of residues from helices TH1-9 and TH5’.

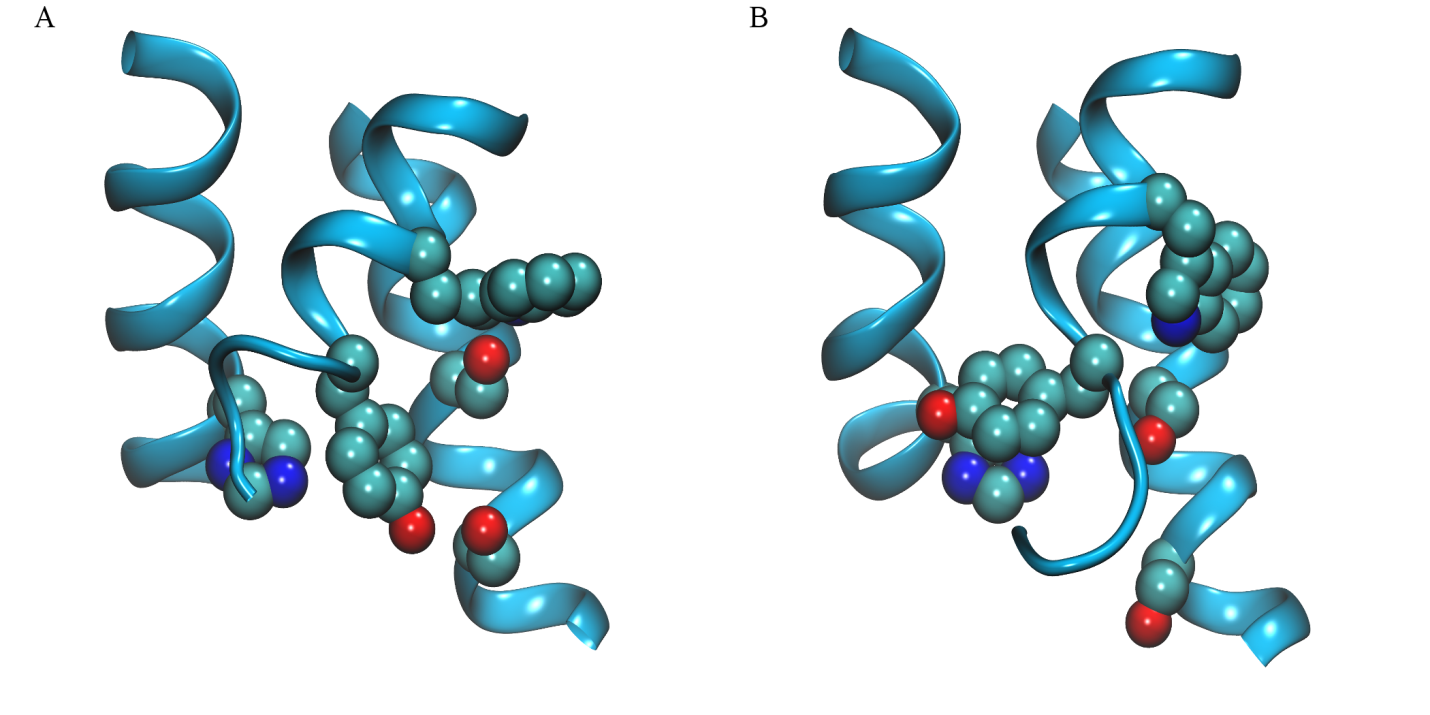

**Figure S12.** (A) Initial and (B) final frames obtained from MD trajectory T3. Side-chains of H251, S332, S336, Y278 and W281 are highlighted in space-filled representation. Oxygen, nitrogen, and carbon atoms are colored in red, blue, and blue, respectively. Helices TH3, TH5, and TH8 are represented in cyan ribbons.

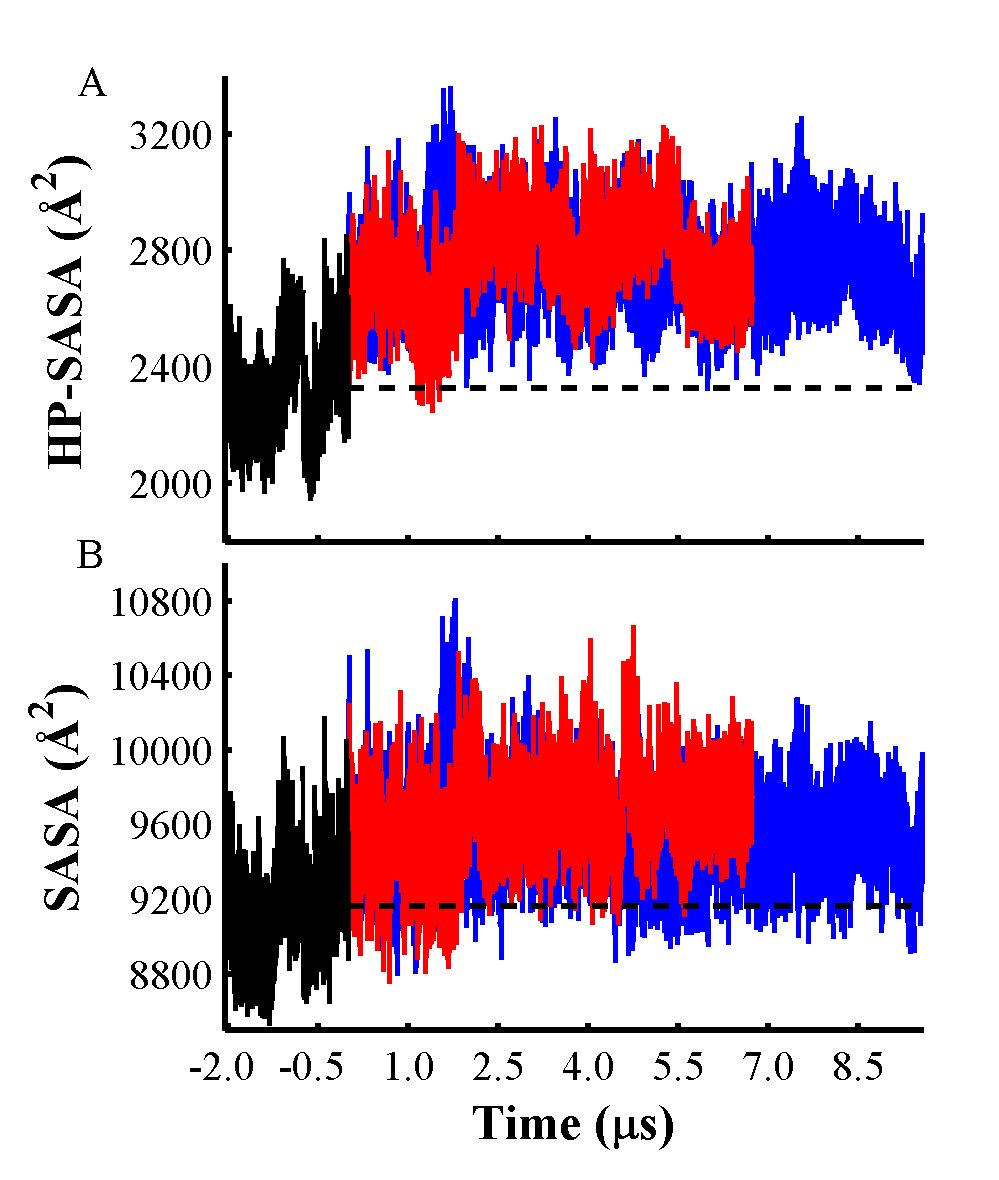

**Figure S13.** SASA traces versus simulation time of hydrophobic and all side-chains in T-domain. (A) Hydrophobic SASA (HP-SASA) of the entire T-domain for T1, T2, and T3, shown in black, red and blue lines, respectively. (B) SASA of all side-chains in T-domain for T1, T2, and T3, shown in black, red and blue lines, respectively. Broken black lines represent the average values of HP-SASA and SASA for T1, 2328 Å^2^ and 9161 Å^2^, respectively.
